# Distinct roles of Nrf1 and Nrf2 in monitoring the reductive stress response to dithiothreitol (DTT)

**DOI:** 10.1101/2022.06.24.497421

**Authors:** Reziyamu Wufur, Zhuo Fan, Jianxin Yuan, Ze Zheng, Shaofan Hu, Guiyin Sun, Yiguo Zhang

## Abstract

Transcription factor Nrf2 (nuclear factor, erythroid 2-like 2, encoded by *Nfe2l2*) has been accepted as a key player in redox regulatory responses to oxidative or reductive stresses. However, it is less or not known about the potential role for Nrf1 (nuclear factor, erythroid 2-like 1, encoded by *Nfe2l1*) in the redox responses, particularly to reductive stress, albeit this ‘fossil-like’ factor is indispensable for cell homeostasis and organ integrity during life process. Here, we examine distinct roles of Nrf1 and Nrf2 in monitoring the defense response to 1,4–dithiothreitol (DTT, serving as a reductive stressor), concomitantly with unfolded protein response being induced by this chemical (also as an endoplasmic reticulum stressor). The results revealed that intracellular reactive oxygen species (ROS) were modestly increased in DTT-treated wild-type (*WT*) and *Nrf1α*^*–/–*^ cell lines, but almost unaltered in *Nrf2*^*–/–ΔTA*^ or *caNrf2*^*ΔN*^ cell lines (with a genetic loss of its transactivation or N-terminal Keap1-binding domains, respectively). This chemical treatment also enabled the rate of oxidized to reduced glutathione (i.e., GSSG to GSH) to be amplified in *WT* and *Nrf2*^*–/–ΔTA*^ cells, but diminished in *Nrf1α*^*–/–*^ cells, along with no changes in *caNrf2*^*ΔN*^ cells. Consequently, *Nrf1α*^*–/–*^, but not *Nrf2*^*–/–ΔTA*^ or *caNrf2*^*ΔN*^, cell viability was reinforced by DTT against its cytotoxicity, as accompanied by decreased apoptosis. Further experiments unraveled that Nrf1 and Nrf2 differentially, and also synergistically, regulated DTT-inducible expression of critical genes for defending redox stress and endoplasmic reticulum stress. In addition, we have also identified that Cys342 and Cys640 of Nrf1 (as redox-sensing sites within its N-glycodomain and DNA-binding domain, respectively) are required for its protein stability and transcription activity.

## 1. Introduction

In order to maintain cell redox homeostasis and organ integrity during healthy life process, almost all cellular life forms have evolutionally established a set of proper defense mechanisms in response to a variety of challengeable (oxidative or reductive) stresses. Those substances such as scavengers, blockers and repair agents of free radicals and reactive oxygen species (ROS) are collectively named antioxidants. Of note, not all reductants have antioxidant properties, because only those that can scavenge free radicals and ROS are *de facto* antioxidants. Recently, thiol compounds have been proved to protect the mitochondria from oxidative stress and relevant damage, which is attributed to their ability to scavenge oxygen and nitrogen free radicals (1) (2) (3). Among them, *1,4-dithiothreitol* (DTT) is a well-known compound, because its structure is similar to that of reduced glutathione (GSH), so that it is often used to preserve the reductive state of sulfhydryl (-SH) groups in many proteins and assist in stabilizing their functional resilience. In fact, it is plausible that a reducing agent with antioxidant properties prevents intracellular inflammatory response (4). DTT can prevent the oxidation of protein cysteine residues, but also interferes with the catalytic role of disulfide isomerase in disulfide bond formation of proteins. From this, DTT is hence reasoned as a stimulator of endoplasmic reticulum (ER) stress (5). Additional ROS, including DTT-relevant free radicals, are also produced to certain degrees. For instance, molecular oxygen is reduced to superoxide free radicals and hydrogen peroxide. Particularly, In the presence of trace metals (e.g., copper and iron), it is reduced to hydroxyl free radicals, resulting in DNA damage and cell apoptosis (6). Such a paradoxical phenomenon, which not only has antioxidant properties, but also can induce reducing stress only when it is excessive, is worth exploring for the in-depth study of DTT(7). Importantly, although DTT has been reported as a treatment-oriented method for some syndromes and complications (8) (9), there are few studies on its application in liver cancer.

In this ever-challenging oxygenated environments, distinct types of cells have evolutionarily armed to adapt to oxidative or reductive stresses in the short term by metabolic reprogramming and in the longer term by genetic reprogramming. The adaptive reprogrammings are determined predominantly by those specified transcription factors-regulated gene expression networks at distinct layers, so as to give rise to a variety of stress responses to adverse external pressures or excessive demands (10,11). Within such multi-hierarchical regulatory networks, the Cap’n’collar (CNC) basic region-Leucine zipper (bZIP) family of transcription factors plays a vital role in maintaining robust redox homeostasis in the internal environments. Two principal CNC-bZIP family members Nrf1 (nuclear factor, erythroid 2-like 2, encoded by *Nfe2l1*) and Nrf2 (encoded by *Nfe2l2*) enable the cellular resistance to challenging redox stresses by mediating proper expression of those cytoprotective genes driven by antioxidant/electrophile response elements (AREs/EpREs) in their promoter regions. Rather, although both are highly homologous, distinctions in their structures and subcellular locations of between Nrf1 and Nrf2 determine potential differences and overlaps in their physiobiological functions, in which such an inter-regulatory relationship of their ‘opposition and unity’ has been existing (12).

In this field, Nrf2 has been preferentially accepted as a master regulator of those antioxidant, detoxification, and cytoprotective genes in response to oxidative or reductive stresses (11,13-16), albeit as a matter of fact that Nrf1, rather than Nrf2, is indispensable for cell homeostasis and organ integrity during normal development and growth, as well as adult life process (10). This is possibly owing to the acute emergent response mediated by transcriptional activity of Nrf2 that is negatively regulated by a thiol-enriched electrophile sensor Keap1 (Kelch-like ECH-associated protein 1), which is attributable to their direct interaction enabling this water-soluble Nrf2 factor to be sequestered within the cytoplasmic subcellular compartments and targeted for its ubiquitination by Cullin 3-based E3 ubiquitin ligase, leading to its protein turnover by the ubiquitin-proteasomal degradation pathway. Only upon stimulation by redox electrophile or nucleophile stress, this thiol-based Keap1 is activated so as to allow Nrf2 to be released and then translocated into the nucleus, resulting in transactivation of target genes. By contrast, the membrane-bound Nrf1 is highly conserved as ‘a living fossil’ of organismal evolution from marine bacteria to mammals (17). The nature-selected Nrf1 possesses a unique intrinsic characteristic to fulfill its special indispensable physiobiological functions that cannot be compensated by Nrf2 and other homologous factors (10). It is of crucial significance to note the fact that *Nrf1*^*–/–*^ cell lines and relevant model animals have suffered from severe endogenous oxidative stress, as manifested by obvious pathological phenotypes, one of which are exemplified by resembling human non-alcoholic steatohepatitis (NASH) and ultimate malignance to hepatoma. This is a rare case of the few typical cell-autonomous defects resulted from spontaneous oxidative stress, as far as we know in the current literature (10,13).

The unique functioning of Nrf1 is dictated by its original transmembrane-topobiology, with specific post-translational modification and selective proteolytic processing during dynamic dislocation from the endoplasmic reticulum (ER) across membranes to enter extra-ER cyto/nucleoplasmic subcellular compartments before gaining access to its cognate genes (10). Thereby, Nrf1 has been identified as an important ER sensor for changes in the intracellular redox, glucose, protein and lipid (including cholesterol) (18,19). Our previous work demonstrated differential and integral contributions of Nrf1 and Nrf2 to coordinately mediating distinct responsive gene expression profiles to the ER stressor tunicamycin (TU) (19) and pro-oxidative stressor *tert*-butylhydroquinone (*t*BHQ)(20). Recently, Nrf1, rather than Nrf2, is further identified as an indispensable redox-determining factor for mitochondrial homeostasis, in addition to the ER-associated proteostasis, by integrating multi-hierarchical responsive signaling pathways through those nuclearly-located respiratory gene controls towards mitochondrially-located gene regulatory networks (21). Herein, we found distinctive roles of Nrf1 and Nrf2 in synergistically monitoring the defense response to DTT as a reductive stressor, concomitantly with unfolded protein response being induced by this chemical (also serving as an ER stressor). Further evidence has been presented, revealing that intracellular reactive oxygen species (ROS) were modestly increased in DTT-treated wild-type (*WT*) and *Nrf1α*^*–/–*^ cell lines, but almost unaltered in *Nrf2*^*–/–ΔTA*^ or *caNrf2*^*ΔN*^ cell lines (with a genetic loss of its transactivation or N-terminal Keap1-binding domains, respectively). This treatment also enabled the rate of oxidized to reduced glutathione (i.e., GSSG to GSH) to be amplified in *WT* and *Nrf2*^*–/–ΔTA*^ cells, but substantially diminished in *Nrf1α*^*–/–*^ cells, along with no changes in *caNrf2*^*ΔN*^ cells. This finding indicates that a potential reductive stress is also induced, possibly by aberrant accumulation of hyperactive Nrf2 in *Nrf1α*^*–/–*^ cells, apart from its severe endogenous oxidative stress. Consequently, *Nrf1α*^*–/–*^, but not *Nrf2*^*–/–ΔTA*^ or *caNrf2*^*ΔN*^, cell viability was reinforced by DTT against its cytotoxicity, as accompanied by decreased apoptosis. Lastly, we have also identified that Cys342 and Cys640 of Nrf1 (as redox-sensing sites situated within its N-glycosylated transactivation domain and its DNA-binding basic region, respectively) are required for this CNC-bZIP protein stability and its transcription activity.

## 2. Materials and methods

### 2.1 Experimental cell lines and culture

The human hepatocellular carcinoma HepG2 cells (i.e., *WT*) were obtained from the American Type Culture Collection (ATCC, Manassas, VA, USA). Three HepG2-derived knockout cell lines *Nrf1α*^*–/–*^, *Nrf2*^*–/–ΔTA*^, and *caNrf2*^*ΔN*^ (with constitutive activation of *Nrf2*) were established in our laboratory as previously described by Qiu *et al* (22). Notably, the fidelity of HepG2 cells had been conformed to be true by its authentication profiling and STR (short term repeat) typing map (Shanghai Biowing Applied Biotechnology Co., Ltd, Shanghai, China). All these cells were incubated in a 37 °C with 5% CO_2_, and allowed for growth in DMEM supplemented with 25 mmol/L glucose (Gibco, USA), 10%(v/v) FBS (Gibco, USA) and 100 units/mL penicillin-streptomycin (MACKLIN, Shanghai, China).

### 2.2 Reagents and antibodies

The chemical *(2S, 3S)-1,4-Bis-sulfanylbutane-2,3-diol* (DTT, CAS No. 3483-12-3) was obtained from Sigma; it is a strong reductant with chemical formula of *C*_*4*_*H*_*10*_*O*_*2*_*S*_*2*_ and *MW* 154.25g/mol. Its reducibility is largely due to the conformational stability after oxidation (containing disulfide bond). In this experiment, 3.09 g DTT powder was completely dissolved in 20 mL 0.01M sodium acetate to obtain 1 M DTT stock solution, which was packed and stored at -20°C before used.

Specific antibody against Nrf1 was made in our own laboratory (23). All nine distinct antibodies against Nrf2 (ab62352), GCLC (ab207777), GCLM (ab126704), HO-1 (ab52947), GPX1 (ab108427), XBP1 (AB109221), ATF4 (ab184909), ATF6 (ab227830) and P4HB (ab137180) were obtained by Abcom (Cambridge, UK). First three antibodies against TALDO (D623398), GSR (D220726) and NQO1 (D26104) were purchased from Sangon Biotech (Shanghai, China). Secondary three antibodies against BIP (bs-1219R), Chop (bs-20669R) and pIRE1 (bs-16698R) were from Bioss (Beijing, China). Third three antibodies against PSMB5 (A1975), PSMB6 (A4054), PSMB7 (A14771) were from ABclonal (Wuhan, China). Lastly, three antibodies to p-eIF2α (#5199) was from CST (Boston, USA), p-PERK (sc-32577) from Santa Crus (CA, USA), and β-actin (TA-09) from ZSGB-BIO (Beijing, China), respectively.

### 2.3 Cell viability assay

All four genotypic cell lines (*WT, Nrf1α*^*–/–*^, *Nrf2*^*–/–ΔTA*^ and *caNrf2*^*ΔN*^) were plated in 96-well plates at a density of 6 × 10^3^ cells per well, each treatment of which was repeated in six separated wells. After the cells completely adhered, they were treated with different concentrations of DTT (0.5, 1.0, 2.0, 3.0 or 4.0 mM) for 24 h or cultured with a single dose of 1 mM DTT for different lengths of time (0, 4, 8, 12, 16, 20 or 24 h). Additionally, MTT reagent (10 ul/well of 10 mg/ml stocked) was used to detect the cell viability.

### 2.4 Intracellular ROS measurement

Equal numbers (3 × 10^5^cells/well in 6-well plate) of experimental cells (*WT, Nrf1α*^*–/–*^, *Nrf2*^*–/–ΔTA*^ and *caNrf2*^*ΔN*^) were allowed for growth 24 h. After reaching 80% of their confluence, the cells were treated by 1 mM DTT for distinct time periods (i.e. 0, 4, 12, 24 h), and then collected. The intracellular ROS were determined by flow cytometry, according to the instruction of ROS assay kit (S0033S, Beyotime, Shanghai, China). Additionally, four different genotypic cells were also plated in 3.5-cm plates with 2.5 × 10^5^ cells/plate and treated as described above. Thereafter the cells were cleaned with a cold PBS buffer, and then incubated with 10 μM of DCFH-DA staining solution (ROS assay kit) at 37 °C for 20 min, before being visualized under a fluorescence microscope.

### 2.5 Assay for the GSSG to GSH ratio

All experimental cells were seeded in 6-well plates (4×10^5^ cells/well). The cells were then treated with 1 mM DTT for different lengths of time, and then collected, before being subjected to determination of the GSSG to GSH ratio according to the instruction of a T-GSH and GSSG assay kit (A061-1-2, Nanjing jiancheng bioengineering institute, Nanjing, China).

### 2.6 The pulse-chase experiments for Nrf1 and its mutants with distinct *trans-*activity

Only four cysteine (Cys) residues of Nrf1 are in its distinct domains: Cys342 situated in its N-glycosylated NST domain, whilst two closer Cys521 and Cys533 are in its Neh6L domain that is highly homologous with Nrf2, and Cys640 is placed in the basic DNA-binding region. They were subjected to site-directed mutagenesis into serine (*S*) residues, respectively. The resulting expression constructs for Nrf1 and its mutants had been transfected into experimental cells, before the cells were treated with 5μg/mL cycloheximide (CHX, a protein synthesis inhibitor) alone or plus the proteasome inhibitor MG132 (10 μM) for different periods of time (0, 0.25, 0.5, 1, 2 or 4 h). Thereafter, these protein samples were collected, and then subjected to measuring their half-lives. In addition, *trans*-activity of Nrf1 and its mutants was determined by the *pGL3-basic-6 × ARE-Luc* reporter assay as described previously (20).

### 2.7 Real-time quantitative PCR analysis of key gene expression

After all experiment cells reached 80% of their confluence, they were treated with 1 mM of DTT for indicated periods of time. Total RNAs were extracted and then subjected to the reaction with a reverse transcriptase to synthesize the first strand of cDNAs. Subsequently, the mRNA levels of examined genes in different cell lines were determined by real-time quantitative PCR (RT-qPCR) with each indicated pairs of their forward and reverse primers (as listed in Table 1). All the experiment were carried out in the Go Taq real-time PCR detection systems by a CFX96 instrument. The resulting data were analyzed by the Bio-Rad CFX Manager 3.1 software.

**Table 1.**
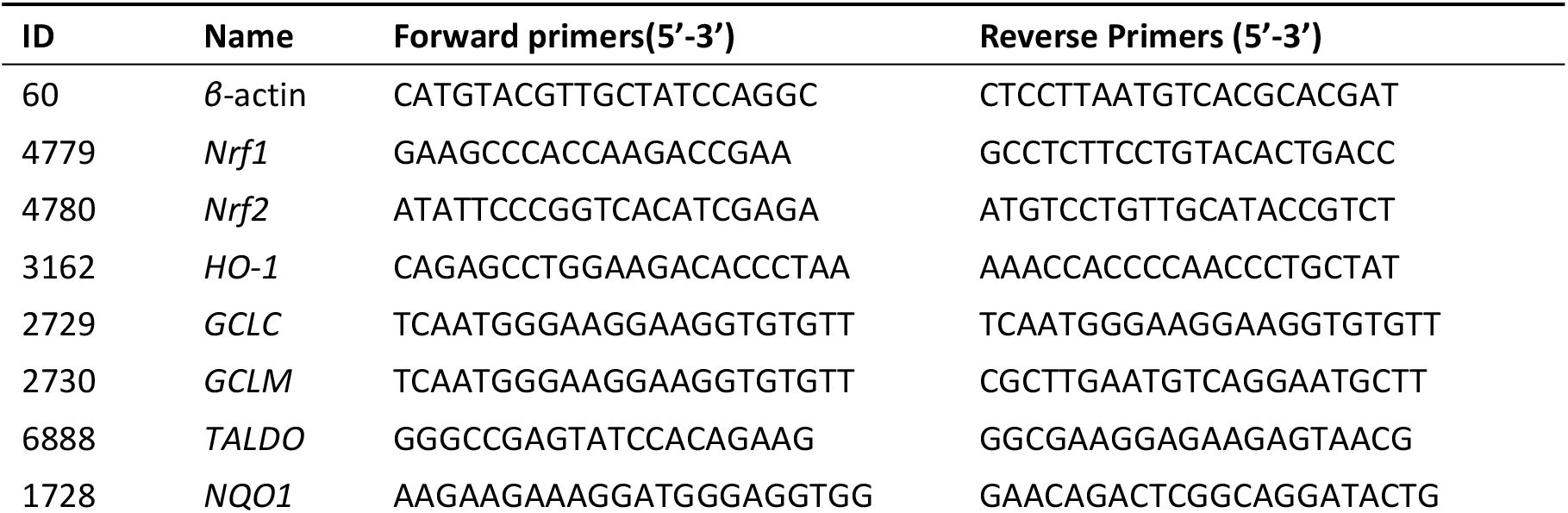
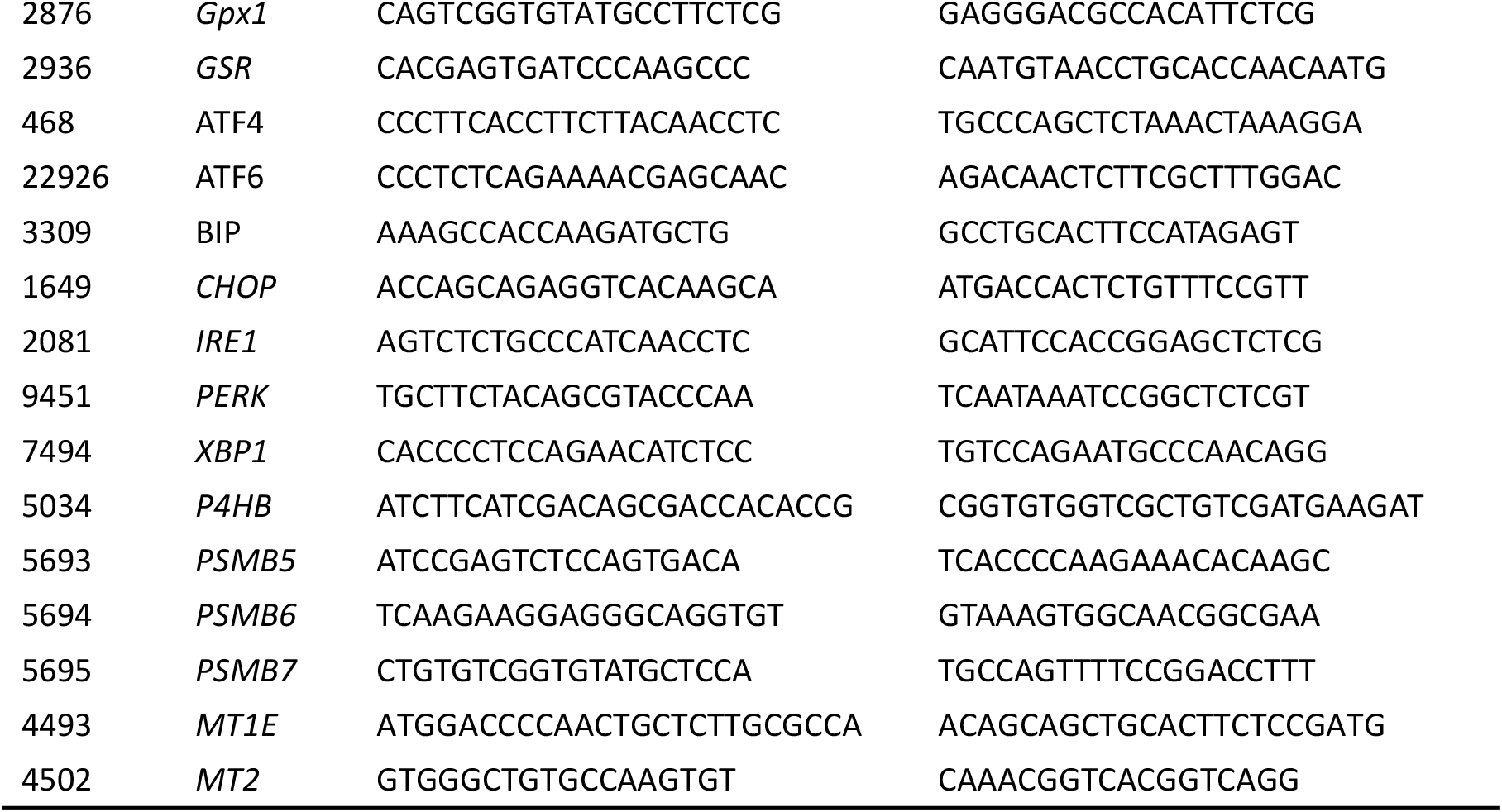
The primer pairs used for q-RT-PCR analysis.

### 2.8 Western blotting analysis of key protein expression

Different genotypic cell lines were plated at 6-well and allowed for an exposure to indicated experimental conditions. The expression abundances of selected proteins were determined by Western blotting. Briefly, after quantitating total proteins in each sample with the BCA protein reagent (P1513-1, ApplyGen, Beijing, China), they were separated by SDS-PAGE gels (8% to12% polyacrylamide) and then transferred on to an PVDF membranes, after blocking in Tris-buffered saline containing 5% non-fat dry milk at room temperature for 1 h, the membranes were incubated with each of specific primary antibodies over night at 4°C. After washing with PBST, the blotted membranes were re-incubated with the secondary antibodies at room temperatures for 2 h. After imaging with Bio-Rad, the intensity of immunoblotted proteins is calculated by the Quantity One software, while β-actin was used as a loading control.

### 2.9 Statistical analysis

The data presented herein are shown as fold changes (*mean±SD*) relative to controls, that were calculated from at least three independent experiments, each of which was performed in triplicates. Statistical significance was assessed by using the one-way ANOVA. And the Tukey’s post hoc test was also used to determine the significance for all pairwise comparisons of interest. These differences between distinct treatments were considered to be statistically significant at *p*<0.05 or *p*<0.01.

## 3. Results

### 3.1 Distinct intervening effects of DTT on different genotypic cell survival, as well on *Nrf1 and Nrf2* expression

Before this experiment, we verified the characteristic protein of four cell lines with different genotypes, and confirmed them to be true as reported previously (19,22) (Fig. 1A). Besides, as mentioned by Xiang *et al* (24), there are four major isoforms derived from human Nrf1α: A and B represent its full-length glycoprotein and deglycoprotein respectively, while C and D denote two distinct lengths of its *N*-terminally-truncated proteins. All these four Nrf1α-derived isoforms were completely deleted upon specific knockout of Nrf1α in *Nrf1α*^*–/–*^ cells (Fig. 1A), but still present in *WT, Nrf2*^*–/–ΔTA*^, and *caNrf2*^*ΔN*^ cell lines. Relatively, the short Nrf1β was obviously decreased in *Nrf1α*^*–/–*^ cells, but significantly augmented in both *Nrf2*^*–/–ΔTA*^, and *caNrf2*^*ΔN*^ cell lines when compared to its equivalent in *WT* cells, indicating that basal abundances of Nrf1 and/or its processing may monitored by Nrf2, besides itself. It is worth noting that Nrf2 was highly expressed in *Nrf1α*^*–/–*^ cell lines, but completely abolished in *Nrf2*^*–/–ΔTA*^ cells. Overall, it is inferable that loss of Nrf1α or Nrf2 may lead to putative inter-regulatory changes in cognate gene expression profiles among these four cell lines, which were used in subsequent experiments.

**Figure 1.**
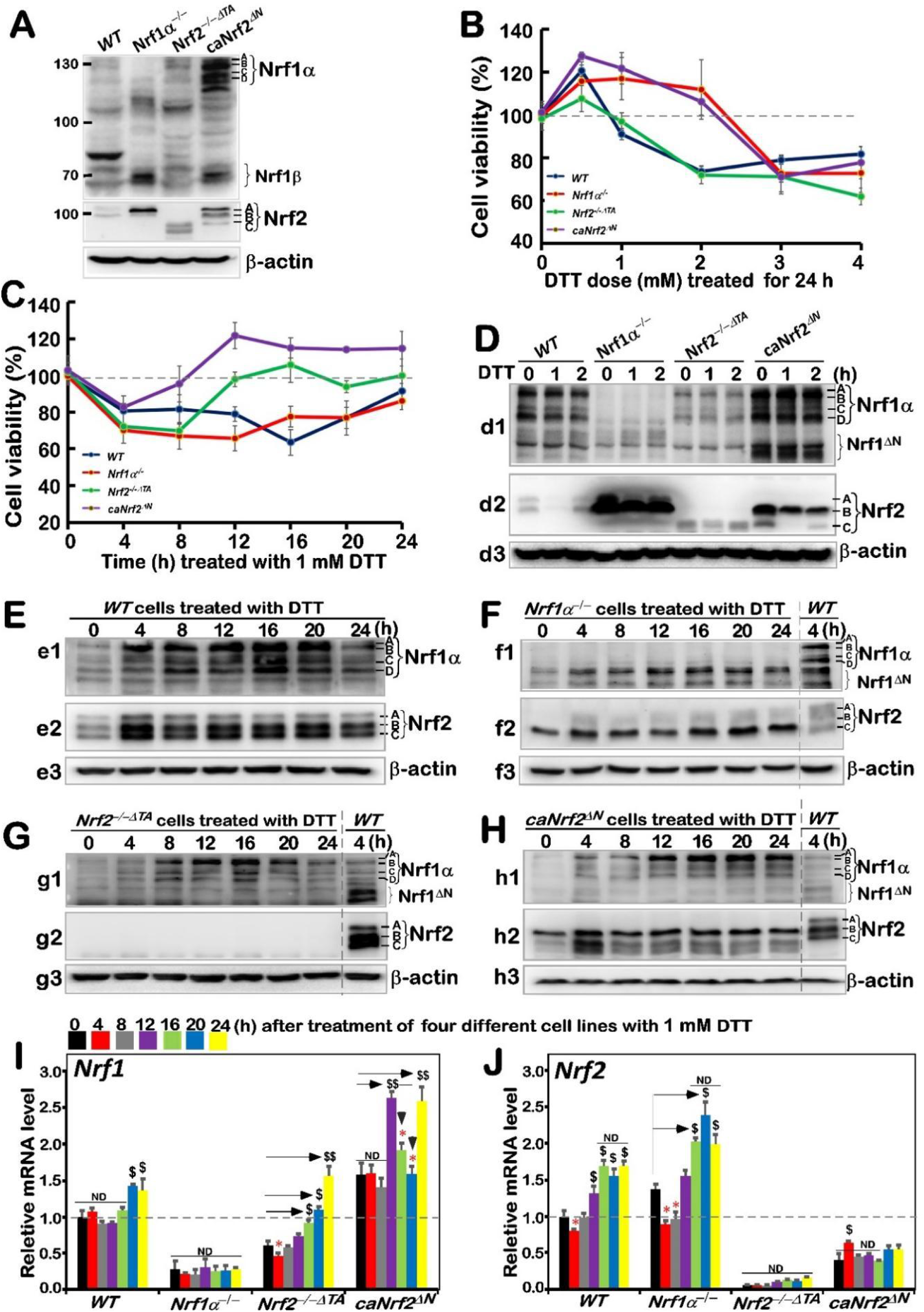
An internal regulatory relationship between Nrf1 and Nrf2 in mediating reductive stress response to DTT. **(A)** Distinct protein levels of Nrf1 and Nrf2 in four different genotypic cell lines *WT, Nrf1α*^*–/–*^, *Nrf2*^*–/–△TA*^ and *caNrf2*^*△N*^ were determined by Western blotting with their specific antibodies. **(B**,**C)** The viability of these cell lines was detected by the MTT-based assay, after they had been treated with different concentrations of DTT for 24 h **(B)** or treated with 1 mM of DTT for different lengths of time **(C). (D to H)** The Nrf1/2 protein expression levels in four genotypic experimental cell lines intervention with 1 mM of DTT for short times (*i*.*e*. 0, 1, 2 h) **(D)** long times (*i*.*e*. 0, 4, 8, 12, 16, 20, 24 h) **(E to H). (I**,**J)** Both *Nrf1* and *Nrf2* mRNA expression levels in different cell lines that has been treated with 1 mM DTT for distinct time periods (*i*.*e*. 0, 4, 8, 12, 16, 20, 24 h) were examined by real-time qPCR. The data are representative of at least three independent experiments being each performed in triplicates. Significant increases (*$, p < 0*.*05; $$, p < 0*.*01*) and significant decreases (*\* p < 0*.*05; ** p < 0*.*01*), in addition to the non-significance (ND), were statistically determined when compared with the corresponding controls (measured at 0 h), respectively. All those protein-blotted bands were also qualified by Quantity One 4.5.2 software as showed in **Figure S1**.

Based on redox characteristics, a reducing compound DTT was employed to explore the effect of foreign substances on cell viability of different genotypes by MTT assay, so to evaluate the formation of formazan precipitates with succinate dehydrogenase in the mitochondria of all living cells only. The result revealed that the viability of all cell lines decreased when they were intervened for 24 h by higher concentrations (3 to 4 mM) of DTT (Fig. 1B), although lower concentrations (0.5-2 mM) of DTT caused modestly enhanced survival of *Nrf1α*^*–/–*^and *caNrf2*^*ΔN*^ cell lines. By contrast, a significant dose-dependent effect of DTT was manifested in *Nrf2*^*–/–ΔTA*^, similarly to that obtained from *WT* cells. Considering such different survival between cells along with cytotoxicity of DTT, we selected a more appropriate dose at 1 mM of this compound to continue intervention of all experimental cell lines for distinct periods of time from 0 to 24 h (*i*.*e*. 0, 4, 8, 12, 16, 20 or 24 h). As shown in Fig 1C, the viability of all four cell lines was modestly decreased within 8 h of DTT intervention. Such decreased viability of *Nrf1α*^*–/–*^ and *WT* cell lines were maintained before 20 h when they was gradually recovered from intervention. By contrast, the resilience of *Nrf2*^*–/–ΔTA*^ and *caNrf2*^*ΔN*^ cell lines appeared to be strikingly recovered from 8 h to 12 h intervention of DTT, and then reached or exceeded their basal levels, respectively. However, the overall viability of these cell lines was not very significant, and thereby several time periods of DTT intervention were selected with the reference value for follow-up experiments to explore the regulatory differences between Nrf1 and Nrf2 in mediating the cellular reductive stress responses to intervention of this reducing compound.

The short-term (*i*.*e*. 0, 1, 2 h) intervention of DTT did not cause a significant difference in Nrf1 abundances in each of other three cell lines except *Nrf1α*^*–/–*^ (Fig. 1D, *d1*), while Nrf2 protein abundances were only modestly decreased in each of other three cell lines except *Nrf2*^*–/–ΔTA*^ (Fig. 1D, *d2*, albeit it was rather highly expressed in *Nrf1α*^*–/–*^ and *caNrf2*^*ΔN*^ cell lines), when compared with their respective basal levels (obtained at time point 0). Thereby, the time of intervention extended from 4 h to 24 h, so as to gain the time-dependent effect of DTT on Nrf1 and Nrf2, as well on their targets.

The resulting data were illustrated in Fig. 1E, revealing significant increases in Nrf1α-derived isoforms, particularly its glycoprotein-A, with the extending time of DTT stimulation from 4 h to 20 h, by comparison with *WT*_*t0*_ as the vehicle control. DTT-inducible expression of Nrf2 was also increased to a considerably higher level after 4 h of this treatment and so higher expression was maintained to 24 h before stopping experiments. Such changes in Nrf1 and Nrf2 when *WT* cells were exposed to DTT for 4 h (i.e., *WT*_*t4*_) served as the ensuing parallel experimental controls. Next, examinations of DTT-treated *Nrf1c*cells unraveled that albeit this protein itself was completely lost, highly-expressed Nrf2 was further enhanced to much higher levels than that obtained from the *WT*_*t4*_ (Fig. 1F). In *Nrf2*^*–/–ΔTA*^ cells, although basal expression of Nrf1 seemed weaker than that of *WT*_*t4*_, its DTT-inducible expression was substantially augmented (Fig. 1G), specifically after this chemical intervention from 8 h to 20 h, whereas Nrf2 was totally abolished. In *caNrf2*^*ΔN*^ cells, the expression levels of Nrf1 and Nrf2 were also significantly incremented by DTT in a time-dependent manner (Fig. 1H). All relevant immunoblots were quantitatively analyzed as shown in supplemental Fig. S1.

Further examinations by RT-qPCR showed that DTT intervention of *WT* cells only caused a marginally-lagged increase in Nrf1 mRNA expression after 20-h stimulation, while Nrf2 mRNA levels were significantly incremented after only 12-h DTT treatment when compared to its basal levels (Fig. 1I vs 1J). In *Nrf1α*^*–/–*^ cells, Nrf2 mRNA expression was also further augmented, albeit Nrf1 expression was largely abrogated. By contrast, although basal Nrf1 expression in *Nrf2*^*–/–ΔTA*^ cells was prevented, but its DTT-stimulated expression was still enhanced after 16-h of this treatment (Fig. 1I). However, it is, much to our surprise, found that although putative constitutive active protein of Nrf2 was present in *caNrf2*^*ΔN*^ cells, its mRNA expression levels were unaffected by DTT (Fig. 1J), but conversely, striking elevation of Nrf1 mRNA expression occurred after 12-h treatment. Overall, these demonstrate there exists a potential inter-regulatory relationship between Nrf1 and Nrf2, albeit under DTT-leading reductive stress conditions.

### 3.2 Differential expression of *Nrf1*/*2-*mediated redox responsive genes induced by DTT in distinct genotypic cells

Our previous study had shown that tBHQ, as a small molecule antioxidant, stimulates mRNA expression of some *ARE*-dependent genes downstream of Nrf1 and Nrf2 (such as *GCLM, GCLC, HO1*, etc.). Here, we examine whether these ARE-driven genes were affected by intervention of DTT as a reducing agent of sulfhydrylation. As anticipated, RT-qPCR results showed that significant time-dependent increases in DTT-inducible mRNA expression of *GCLM* and *GCLC* encoding two subunits of GSH synthesis rate-limiting enzyme that is of crucial importance in redox process (25) in DTT-treated *WT* cells (Fig. 2A, 2B). Knockout of *Nrf1α*^*–/–*^ cells caused a faster significant increase in DTT-inducible expression *GCLM* from 4-h to its maximum at 16 h, which was then maintained to 24 h before stopping experiments (Fig. 2A), while only a modest increase in *GCLC* expression was apparently lagged to occur from 16 h stimulation by DTT (Fig. 2B). By sharp contrast, *Nrf2*^*–/–ΔTA*^ cells only displayed a marginal increase in expression of *GCLM*, whereas *GCLC* was unaffected by DTT (Fig. 2A *vs* 2B). Conversely, such DTT-induced expression of *GCLM* was completely abolished in *caNrf2*^*ΔN*^ cells, albeit its basal expression was enhanced, while significantly inducible expression of *GCLC* was greatly lagged until 20 h to 24 h treatment. Collectively, these imply that differential expression of *GCLM* and *GCLC* is monitored by inter-regulated Nrf1 and Nrf2 in a time-dependent fashion.

**Figure 2.**
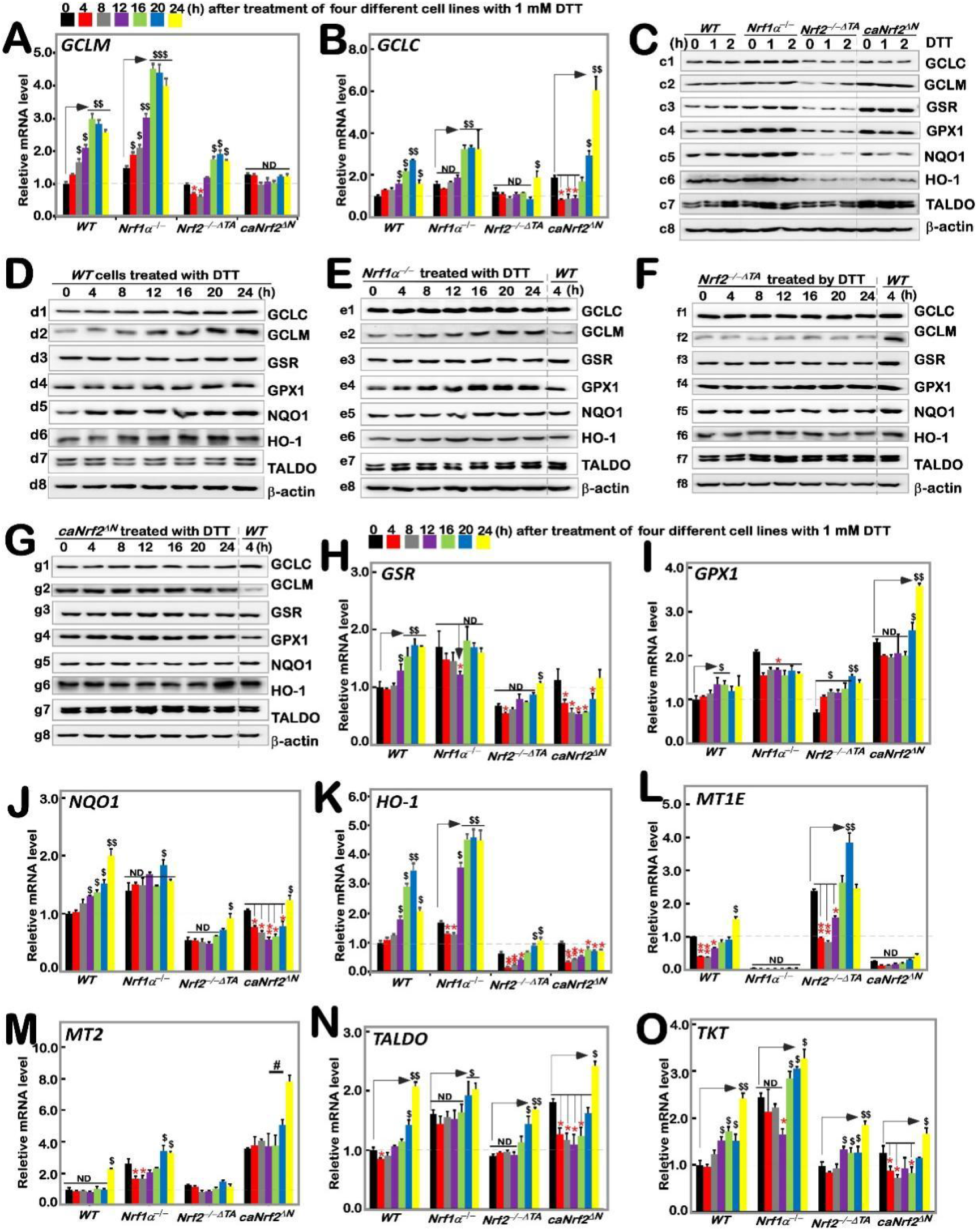
Distinct time-dependent expression of Nrf1/2-mediated redox response genes to DTT in different cell lines. Different genotypic cell lines *WT, Nrf1α*^*–/–*^, *Nrf2*^*–/–△TA*^ and *caNrf2*^*△N*^ were or were not treated with 1 mM DTT for 0 to 24 h, before both basal and DTT-inducible mRNA levels of all examined genes were determined by RT-qPCR. These genes included GCLM **(A)**, GCLC **(B)**, GSR **(H)**, GPX1 **(I)**, NQO1 **(J)**, HO-1 **(K)**, MT1E **(L)**, MT2 **(M)**, TALDO **(N)** and TKT **(O)**. The resulting data were shown as fold changes (*mean ± SD, n = 3 × 3*), which are representative of at least three independent experiments being each performed in triplicates. Significant increases (*$, p < 0*.*05; $$, p < 0*.*01*) and significant decreases (*\* p < 0*.*05; ** p < 0*.*01*), in addition to the non-significance (ND), were statistically analyzed when compared with their corresponding controls (measured at 0 h), respectively. These experimental cells were or were not treated with 1 mM for short times (*i*.*e*. 0, 1, 2 h) **(C)** or long times (*i*.*e*. 0, 4, 8, 12, 16, 20, 24 h), before basal and DTT-inducible protein abundances of GCLC (*c1 to g1*), GCLM (*c2 to g2*), GSR (*c3, d3 to g3*), GPX1 (*c4 to g4*), NQO1 (*c5 to g5*), HO-1 (*c6 to g6*) and TALDO (*c7 to g7*) were determined by Western blotting with indicated antibodies, whilst *β*-actin served as a loading control. The intensity of those immunoblots was also quantified by the Quantity One 4.5.2 software as showed in **Figure S2**.

Taking *WT* (at *t*_*0*_) as the control, basal protein abundances of GCLC and GCLM were upregulated in *Nrf1α*^*–/–*^ and *caNrf2*^*ΔN*^ cell lines, but significantly down-regulated in *Nrf2*^*–/–ΔTA*^ cells (Fig. 2C, *c1 & c2*). However, no significant changes or even decreases in DTT-inducible GCLC and GCLM proteins for shorter-term of 1 h to 2 h were determined all examined cell lines (Fig. 2C and Fig. S2). Upon DTT intervention of *WT* cells for the longer-terms, significant changes in GCLC and GCLM proteins were observed to increment with the increasing time of administration (Fig. 2D, *d1,d2*). After knockout of *Nrf1α*^*–/–*^, GCLM changed rather significantly rather than that GCLC did, with a time-dependent gradual increase (Fig. 2E, *e1,e2*); Knockout of *Nrf2*^*–/–ΔTA*^ enabled *GCLM* expression to become fainter, while GCLC appeared to be unaltered by comparison to the *WT* (at *t*_*4h*_) control (Fig. 2F, *f1,f2*). Besides, *caNrf2*^*ΔN*^ cells had no further stimulated increases in GCLC and GCLM in response to DTT, albeit its basal levels were higher than that of the *WT* (at *t*_*4h*_) (Fig. 2G, *g1,g2*).

As mentioned herein, the oxidized glutathione disulfide (GSSG) can be reduced to GSH form by glutathione-disulfide reductase (GSR, a central enzyme of antioxidant defense), in this redox cycle where glutathione peroxidase 1 (GPX1) can also achieve the purpose of oxidative detoxification in cells by reducing some peroxides. When compared with *WT* (at *t*_*0*_) cells, basal mRNA levels of *GSR* and *GPX1* were evidently up-expressed in *Nrf1α*^*–/–*^ and *caNrf2*^*ΔN*^ cell lines, but rather substantially down-expressed in *Nrf2*^*–/–ΔTA*^ cells (Fig. 2H, 2I). After DTT treatment, transcriptional expression of *GSR* in *WT* cells was significantly up-regulated in a time-dependent manner from 12 h, whereas inducible expression of *GPX1* was modestly upregulated by this chemical. Intriguingly, a DTT-inducible decrease, rather than increase, of *GSR* occurred only at 12 h treatment of *Nrf1α*^*–/–*^ cells (Fig. 2H), as accompanied by obvious down-regulation of *GPX1* (Fig. 2I). *Nrf2*^*–/–ΔTA*^ cells only displayed a lagged increase in mRNA expression of *GSR* at 24 h stimulation (Fig. 2H), but biphasic induction of *GPX1* by DTT occurred respectively at 4 h and 20 h, albeit its basal levels were lowered (Fig. 2I). Conversely, *caNrf2*^*ΔN*^ cells exhibited an obvious time-dependent DTT-inducible decrease of *GSR*, along with lagged induction of *GPX1* by this treatment for 20 h to 24 h, which was rather significantly higher than its basal levels (Fig. 2I). Further examinations of GSR and GPX1 proteins revealed no significant changes in their inducible expression upon short-term intervention of all examined cells by DTT, although their basal levels were highly up-expressed in *Nrf1α*^*–/–*^ and *caNrf2*^*ΔN*^ cell lines, but down-regulated in *Nrf2*^*–/–ΔTA*^ cells by comparison to the *WT* (at *t*_*0*_) control (Fig. 2C, *c3,c4* and Fig. S2). By continuing to detect the longer-term DTT intervening effects on *WT* cells, it were observed almost no significantly inducible changes in GSR and GPX1 abundance, except from a marginal induction being lagged at 24 h (Fig. 2D, *d3,d4*). Similarly to the control *WT* (at *t*_*4*_), almost no induction of GSR by DTT were also observed in *Nrf1α*^*–/–*^ cells, but as compared by enhanced expression of GPX1 in a time-dependent manner (Fig 2E, *e3,e4*). In *Nrf2*^*–/–ΔTA*^ cells, considerably fainter expression of GSR was not induced by DTT, while GPX1 was also not triggered by this drug (Fig 2F, *f3,f4*). However, *caNrf2*^*ΔN*^ cells displayed a modest stimulated expression of GPX1, but not GSR in response to DTT (Fig 2G, *g3,g4*). Together, such distinct expression profiles of GSR and GPX1 at mRNA and protein levels may be attributable to coordinated inter-regulation by Nrf1 and Nrf2.

The expression of downstream genes *NQO1* and *HO-1* closely related to *Nrf1* and *Nrf2* was also increased with time-dependent induction of DTT in *WT* cells (Fig. 2J and 2K). When compared with the *WT*_*t0*_ control, basal mRNA expression levels of *NQO1* and *HO-1* were upregulated in *Nrf1α*^*–/–*^ cells, but down-regulated in *Nrf2*^*–/–ΔTA*^ cells, along with almost no changes in *caNrf2*^*ΔN*^ cells. The inducible expression of *HO-1* was further augmented by DTT stimulation of *Nrf1α*^*–/–*^ cells for 12 h to 24 h (Fig. 2K), whereas *NQO1* was largely unaffected by this chemical, except form a marginal increase lagged at 20 h (Fig. 2J). Intriguingly, *NQO1* and *HO-1* were roughly unaltered or down-regulated by DTT, except from a slightly stimulated expression lagged at 24 h of this treatment in *Nrf2*^*–/–ΔTA*^ and *caNrf2*^*ΔN*^ cell lines. Further investigation of NQO1 and HO-1 revealed a largely similar trend of changes in their protein expression to that of mRNAs. By comparison to *WT* (at *t*_*0*_), basal NQO1 and HO1 expression abundances were enhanced in *Nrf1α*^*–/–*^ cells, but suppressed in either *Nrf2*^*–/–ΔTA*^ or *caNrf2*^*ΔN*^ cells (Fig. 2C, *c5,c6*). Short term DTT intervention of examined cells for 1 h to 2 h also unraveled no significant changes or even a modest decreased trend in NQO1 and HO1 proteins. However, long-term intervention of *WT* cells with DTT from 4 h to 24 h led to gradually incremented abundances of NQO1 and HO-1 in a time-dependent fashion (Fig 2D, *d5,d6*). Further comparison with the *WT (*at *t*_*4*_) control indicated that DTT intervention of *Nrf1α*^*–/–*^ cells for 4 h to 24 h also gave a modest gradual increased expression trend of NQO1, while HO-1 reached a relative inducible expression peak at 12 h and then gradually weakened (Fig 2E, *e5,e6*). In *Nrf2*^*–/–ΔTA*^ cells, NQO1 and HO-1 were not only weaker than those in *WT (*at *t*_*4*_), but also not induced by DTT (Fig 2F, *f5,f6*). Similarly, *caNrf2*^*ΔN*^ cells also manifested no significant changes in NQO1 and HO-1 in response to DTT, with an exception of only slight HO-1 induction at 24 h (Fig 2G, *g5,g*6).

Herein, we also examined the expression of *TALDO* (encoding a key enzyme in the non-oxidative pentose phosphate pathway to yield NADPH so as to maintain a reduced state of glutathione, thus protecting cells from oxygen free radicals (26)), *TKT* (encoding thiamine dependent enzyme to guide excess phosphate sugar to glycolysis in the pentose phosphate pathway), *MT1E* and *MT2* (two members the metal sulfur family that act as antioxidants in the steady-state control of metals in cells and prevent the production of hydroxyl radicals (27)). As shown in, biphasic changes in *MT1E* mRNA expression were observed in *WT* cells, which was first decreased, then gradually recovered and even increased as the time of DTT intervention was extended to 24 h of its maximum response. Loss of *Nrf1α*^*–/–*^ led to a complete abolishment of basal and DTT-inducible mRNA expression of *MT1E* (Fig. 2L). Intriguingly, similar results were obtained from *caNrf2*^*ΔN*^ cells. Conversely, loss of *Nrf2*^*–/–ΔTA*^ led to a substantial increase in basal *MT1E* expression, but its DTT-inducible expression changes were triphasic, which was first inhibited from 4 h to 12 h, then recovered at 16 h and induced to the maximum at 20 h, but finally returned to its basal level (Fig 2L). For *MT2*, only a lagged DTT-stimulated increase occurred at 24 h treatment of *WT* cells (Fig. 2M). When compared with the *WT*_*t0*_ control, basal MT2 expression was significantly up-regulated in *Nrf1α*^*–/–*^ and *caNrf2*^*ΔN*^ cell lines. A biphasic change in its DTT-inducible expression was exhibited in *Nrf1α*^*–/–*^ cells. Similarly lagged DTT-triggering expression of MT2 was also observed in *caNrf2*^*ΔN*^ cells. Conversely, *Nrf2*^*–/–ΔTA*^ cells gave a marginal increase of basal MT2 levels, its DTT-stimulated expression was completely abolished (Fig. 2M). Further examination unraveled that basal mRNA expression levels of *TALDO* and *TKT* were up-regulated in *Nrf1α*^*–/–*^and *caNrf2*^*ΔN*^, rather than *Nrf2*^*–/–ΔTA*^, cell lines, when compared with the *WT*_*t0*_ control (Fig. 2N, 2O). Similar changes in *TALDO* protein levels were obtained (Fig. 2C). Interestingly, significant increases in *TALDO* and *TKT* expression levels were stimulated by DTT in *WT* cells. Knockout of *Nrf1α*^*–/–*^ or *Nrf2*^*–/–ΔTA*^ still gave rise to lagged induction of *TALDO* and *TKT* within 16 h to 24 h after DTT intervention (Fig. 2N, 2O), while *caNrf2*^*ΔN*^ displayed a biphasic change in *TALDO and TKT* expression levels that was first decreased, and then recovered and increased during DTT stimulation with an inducible maximum occurring at 24 h) (Fig 2N, 2O). However, a little or no effects of DTT on the TALDO protein expression in all examined cells (Fig. 2C to 2G).

### 3.3. Differential requirements of *Nrf1* and *Nrf2* for the ER stress responsive genes stimulated by DTT

It was reported that Nrf2 is significantly up-regulated by aggregated β-amyloid-mediated ER stress and activated by PERK as a canonic ER stress sensor (28,29). Our previous work had shown differential and integral roles of Nrf1 and Nrf2 in mediating the unfolded protein response (i.e., UPR^ER^) to the classic ER stressor tunicamycin (19). Herein, we examined whether Nrf1 and Nrf2 are required for monitoring putative ER stress response to DTT, which interferes disulfide bond formation during protein folding towards maturation. Thus, it is reasoned that one of the first targets of oxidative protein folding attacked by DTT should be protein disulfide isomerase (PDI, which can also act as a reductase to cut those protein disulfide bonds attached to the cell surface, aside from an ER chaperone to inhibit misfolded protein aggregation (30,31)). As expected, DTT-stimulated mRNA expression levels of *PDI* were up-regulated in all four examined cell lines, even upon loss of *Nrf1α*^*–/–*^ or *Nrf2*^*–/–ΔTA*^, but its basal enhancement occurred in *Nrf1α*^*–/–*^ and *caNrf2*^*ΔN*^, rather than *Nrf2*^*–/–ΔTA*^, cell lines (Fig. 3A). Such a rebound effect on transcriptional expression of *PDI* is likely controlled by a feedback loop coordinated with Nrf1 and Nrf2. However, its protein expression abundances were significantly incremented by DTT in *Nrf1α*^*–/–*^ cells, but only modestly enhanced in other three cell lines (Fig. 3J to 3M). This implies a possibly enhanced stability of PDI may be attributable to *Nrf1α*^*–/–-*^-impaired proteasomal degradation, particularly under reductive stress conditions. Next, it was, to our surprise, found that mRNA expression levels of GRP78 (a pivotal partner with three ER sensors PERK, IRE1 and ATF6) were rapidly increased to its maximum induction by 4-h intervention of DTT in all four examined cell lines, and then gradually decreased to its basal levels or to rather lower extents, as stimulation time was extended to 24 h (Fig. 3B). Similar biphasic changes in GRP78 protein abundance were also determined in these four cell lines (Fig. 3J, *j2* to 3M, *m2*). However, it is notable that the extents of DTT-inducible mRNA and protein expression in *Nrf1α*^*–/–*^ cells were rather lower than those obtained from *WT, Nrf2*^*–/–ΔTA*^ and *caNrf2*^*ΔN*^, cell lines, but almost no differences in induction of GRP78 were observed between *Nrf2*^*–/–ΔTA*^ and *caNrf2*^*ΔN*^ cell lines. This indicates that Nrf1, rather than Nrf2, is required for bidirectional regulation of GRP78 at distinct levels in response to DTT.

**Figure 3.**
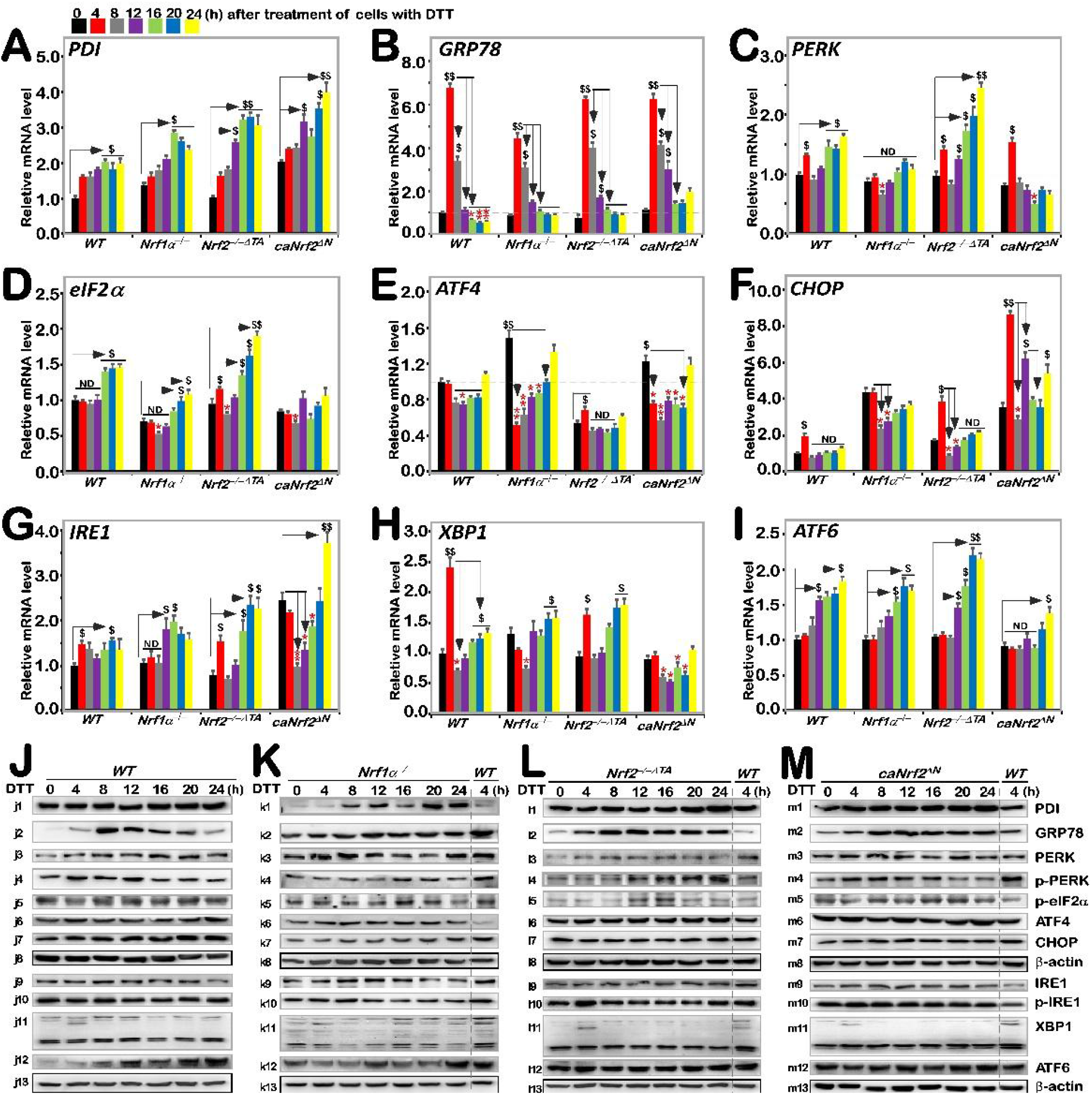
Distinct roles of Nrf1 and Nrf2 in the ER stress response induced by DTT. Four genotypic cell lines of *WT, Nrf1α*^*–/–*^, *Nrf2*^*–/–△TA*^ and *caNrf2*^*△N*^ were or were not treated with 1 mM DTT for 0 to 24 h, before basal and DTT-inducible mRNA expression levels of those ER stress-related genes were determined by RT-qPCR. Those genes included PDI **(A)**, GRP78 **(B)**, PERK **(C)**, eIF2α **(D)**, ATF4 **(E)**, CHOP **(F)**, IRE1 **(G)**, XBP1 **(H)** and ATF6 **(I)**. The data were shown as fold changes (*mean ± SD, n = 3 × 3*), which are representative of at least three independent experiments being each performed in triplicates. Significant increases (*$, p < 0*.*05; $$, p < 0*.*01*) and significant decreases (*\* p < 0*.*05; ** p < 0*.*01*), in addition to the non-significance (ND), were statistically analyzed when compared with their corresponding controls (measured at 0 h). After similar treatment of *WT* **(J)**, *Nrf1α*^*–/––*^ **(K)**, *Nrf2*^*–/–△TA*^ **(L)**, and *caNrf2*^*△N*^ **(M)** cell lines for distinct lengths of time (*i*.*e*. 0, 4, 8, 12, 16, 20, 24 h), those inducible protein changes of PDI (*j1 to m1*), GRP78 (*j2 to m2*), PERK (*j3 to m3*), p-PERK (*j4 to m4*), p-eIF2α (*j5 tom5*), ATF4 (*j6 to m6*), CHOP (*j7 to m7*), IRE1 (*j9 to m9*), p-IRE1 (*j10 to m10*), XBP1 (*j11 to m11*)and ATF6 (*j12 to m12*) were determined by Western blotting with indicated antibodies, whilst *β*-actin served as a loading control. The intensity of those immunoblots, representing different protein expression levels, was also quantified by the Quantity One 4.5.2 software as shown in **Figure. S3**).

By further examination of three ER stress responsive genes, it was revealed that a modest bimodal induction of *PERK* by DTT occurred early at 4 h and later after 16 h of this chemical treatment, respectively, in *WT* cells, and such bimodality was further augmented in *Nrf2*^*–/–ΔTA*^ cells (Fig. 3C), but was substantially blunted in *Nrf1α*^*–/–*^ cells. Of note, the early peak was abolished by knockout of *Nrf1α*, while the latter peak was abrogated and even suppressed by *caNrf2*^*ΔN*^. This implies that Nrf1 and Nrf2 bi-directionally positively and negatively regulate expression of *PERK*, particularly its lagged induction by DTT, respectively. In *WT* cells, phosphorylated protein of p-PERK was significantly stimulated by DTT at 4 h and then decreased to rather lower extents (Fig. 3J, *j4*). A similar, but modest, induction pattern of p-PERK by DTT was manifested in *caNrf2*^*ΔN*^ cells (Fig. 3M, *m4*), but this induction seemed to be abolished by knockout of *Nrf1α*^*–/–*^ (Fig. 3K, *k4*). Conversely, in *Nrf2*^*–/–ΔTA*^ cells, DTT-inducible expression of p-PERK was gradually incremented from 12 h to 24 h of its maximum (Fig. 3L, *l4*). In addition, no significant changes in total PERK were observed in all four examined cell lines, except for partial attenuation by knockout of *Nrf1α*^*–/–*^ within an indicated period of time (Fig. 3J to 3M, and Fig. S3). Collectively, induction of the PERK signaling by DTT is also positively and negatively monitored by Nrf1 and Nrf2, respectively.

The downstream eIF2α-ATF4-CHOP of PERK signaling was also examined herein. The results showed only marginal induction of *eIF2α* and *CHOP* at mRNA expression levels after 16 h or early at 4 h of DTT intervention, respectively, along with no induction or even decreases of *ATF4* in *WT* cells (Fig. 3D to 3F). By contrast, basal and/or DTT-inducible *eIF2α* expression levels were repressed by *Nrf1α*^*–/–*^, but enhanced by *Nrf2*^*–/–ΔTA*^, even though unaffected by *caNrf2*^*ΔN*^ (Fig. 3D). In addition, the phosphorylated eIF2α expression was modestly induced by DTT in *Nrf2*^*–/–ΔTA*^ cells, but no marked changes were in other three cell lines (Fig. 3J to 3M, and Fig. S3). These indicate that Nrf1 and Nrf2 regulate eIF2α by a similar way to monitor PERK expression. However, DTT intervention led to significant down-regulation of *ATF4* in in *Nrf1α*^*–/–*^ and *caNrf2*^*ΔN*^ cell lines when compared with *WT* cells, although upregulation of its basal mRNA expression levels in *Nrf1α*^*–/–*^ and *caNrf2*^*ΔN*^ cells with its suppression in *Nrf2*^*–/–ΔTA*^ cells (Fig. 3E). However, no striking changes in ATF4 protein levels in all examined cells (Fig. 3J to 3M). Hence, it is inferable that Nrf2 may be required for DTT-stimulated trans-repression of *ATF4*. Moreover, only early induction of *CHOP* by DTT occurred, to a lower degree, at 4h treatment of *WT* cells, but also further amplified in *Nrf2*^*–/–ΔTA*^ and *caNrf2*^*ΔN*^, but not in *Nrf1α*^*–/–*^ cell lines, albeit its basal expression was also upregulated in *Nrf1α*^*–/–*^ cells (Fig. 3F). Also, no significant changes in CHOP protein levels were determined in all four cell lines (Fig. 3J to 3M). This implies that Nrf1 is likely required for early induction of *CHOP* by DTT. Overall, these indicate that differential and integral roles of Nrf1 and Nrf2 in monitoring the PERK signaling to its downstream eIF2α-ATF4-CHOP pathway.

A bimodal of the IRE1-XBP1 signaling at their mRNA levels was also induced by DTT in *WT* cells. Such induction of *IRE1* was significantly enhanced in *Nrf2*^*–/–ΔTA*^ cells, though its basal levels were down-regulated (Fig. 3G). The early peak of *IRE1* induction by DTT was abrogated or suppressed in *Nrf1α*^*–/–*^ or *caNrf2*^*ΔN*^ cells, but its later peak was unaffected or augmented in the two cell lines, respectively. The marked early peak of *XBP1* induced by DTT was observed in *WT* cells, decreased by *Nrf2*^*–/–ΔTA*^ and abolished by *Nrf1α*^*–/–*^ or *caNrf2*^*ΔN*^ (Fig. 3H), whereas the secondary peak was also abolished by *caNrf2*^*ΔN*^, but unaffected by *Nrf1α*^*–/–*^ *or Nrf2*^*–/–ΔTA*^. Furthermore, time-dependent increments of *ATF6* mRNA expression occurred from 12 h to 24 h DTT-treated *WT, Nrf1α*^*–/–*^ or *Nrf2*^*–/–ΔTA*^, but not *caNrf2*^*ΔN*^ cells (Fig. 3I). In addition, no obvious changes in IRE1, p-IRE1 and XBP1 proteins were in all four examined cell lines, but ATF6 protein levels were, to different extents, enhanced by DTT in *WT, Nrf1α*^*–/–*^, *caNrf2*^*ΔN*^, rather than *Nrf2*^*–/–ΔTA*^, cell lines (Fig. 3J to 3M). Taken together, these demonstrate that Nrf1 and Nrf2 differentially regulate the ER-stress responsive genes to DTT.

### 3.4 Distinct intracellular redox changes among different genotypic cell lines in response to DTT

As a collective term, ROS include superoxide anion, hydrogen peroxide, and all other oxygenated active substances, but are still hard to be accurately detected, because they have strong oxidation activity to be exerted for a short retention time while they are easy to be removed by antioxidants. Therefore, visualization of ROS by fluorescence microscopy and its quantification by flow cytometry are usually used to detect their changing status evaluated by the green fluorescence raised from the DCFH dye reaction with intracellular ROS, to assess the difference of antioxidant capacity (32). The results showed that basal ROS levels in *Nrf1α*^*–/–*^ cells was obviously higher than that of *WT*_*t0*_, while a slight increase in the yield of ROS was observed after 4 h stimulation by DTT, before being decreased to lower extents than its basal levels (Fig. 4, A to C). Such a slight DTT-stimulated rise in ROS was also observed in *WT* cells before being decreased and then recovered to its basal levels. By contrast, basal ROS levels were also apparently increased in *Nrf2*^*–/–ΔTA*^ cells, but this status seemed to be unaffected by DTT intervention (Fig 4, A to C). Intriguingly, no differences in basal and DTT-stimulated ROS levels in *caNrf2*^*ΔN*^ cells were determined by comparison to *WT* controls. Besides, similar changes in the intensity of fluorescence arising from intracellular DCFH-DA dye observed by microscopy were also fully consistent with the results as described above (Fig 4D). Altogether, these demonstrate that albeit both Nrf1 and Nrf2 are responsible for endogenous antioxidant cytoprotecton, Nrf1 rather than Nrf2 is required for DTT-stimulated antioxidant defense response.

**Figure 4.**
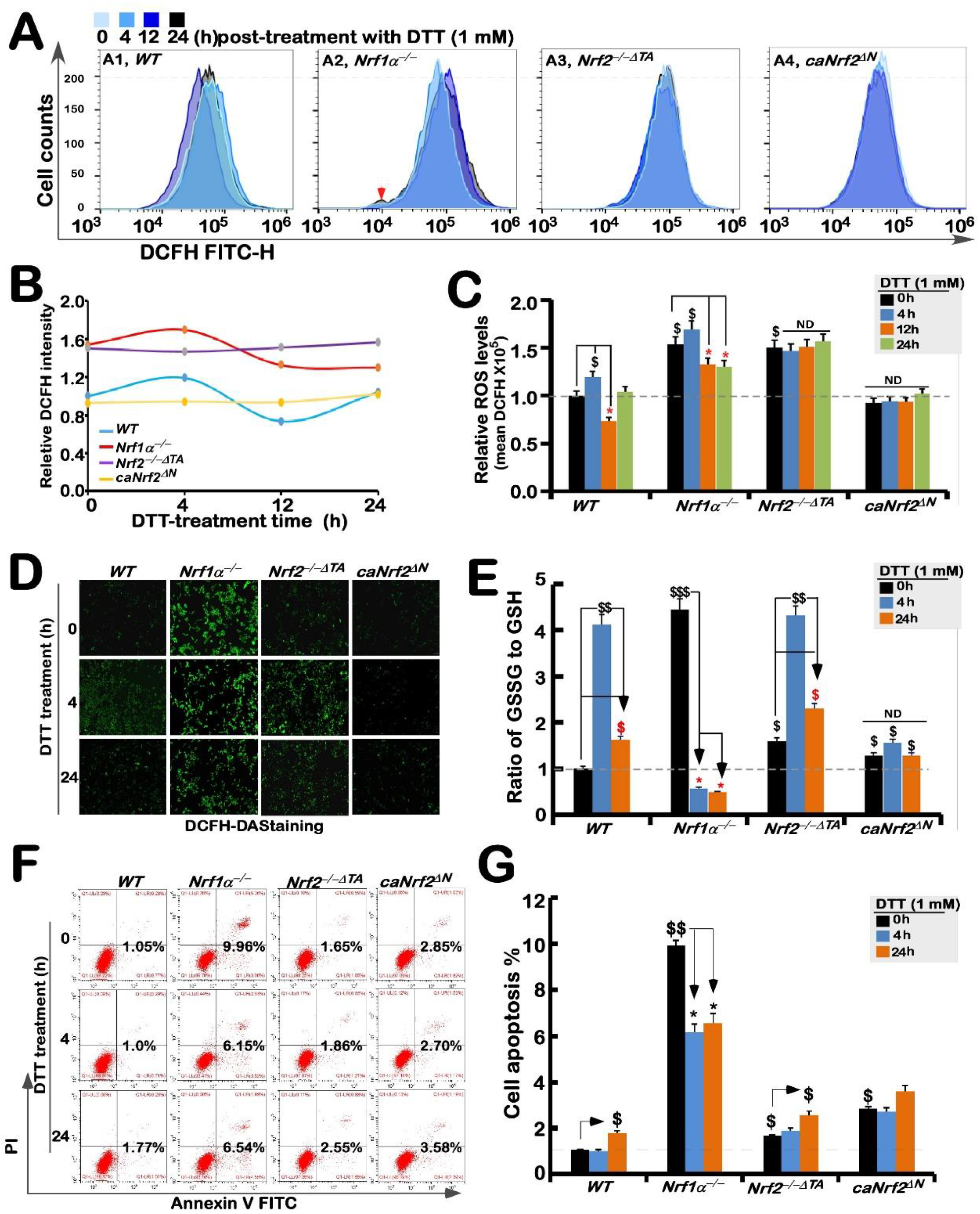
Changes in both ROS levels and ratio of GSSG to GSH are accompanied by distinct apoptosis of different genotypic cell lines against DTT. **(A)** Experimental cell lines *WT, Nrf1α*^*–/–*^, *Nrf2*^*–/–△TA*^ and *caNrf2*^*△N*^ were allowed for treatment with 1 mM DTT for different time periods (i.e., 0, 4, 12 and 24 h), before they were subjected to flow cytometry analysis of their intracellular ROS levels by the DCFH-DA fluorescent intensity. The data of ROS products and differences were further analyzed by FlowJo 7.6.1 software as shown in **(B**,**C). (D)** After DCFH-DA staining, the ROS fluorescent images in different cell lines were obtained under microscope. **(E)** The intracellular GSH and GSSH levels were measured and also repeated three times, each of which was performed in triplicates. Statistic significances were calculated: *$$, p <0*.*01*, and *$, p < 0*.*05* indicate significant increases by comparing each basal value of [*Nrf1α*^*–/–*^]_*T0*_, [*Nrf2*^*–/–△TA*^]_*T0*_, and [*caNrf2*^*ΔN*^]_*T0*_ with that of [*WT*]_*T0*_, while both *\* p < 0*.*05* and *\*\* p < 0*.*01* denote significant decreases of those values from each of cell lines treated by DTT for 4 h (*i*.*e*., [X]_*T4*_) or 24 h (*i*.*e*., [X]_*T24*_) versus its untreated [X]_*T0*_ value in the same groups. **(F)** Distinct cell lines of *WT, Nrf1α*^*–/–*^, *Nrf2*^*–/–△TA*^ and *caNrf2*^*△N*^ were (or were not treated) with 1 mM DTT for different lengths of time. Subsequently, the cells were incubated with a binding buffer containing Annexin V-FITC and propidium iodide (PI) for 15 min, before being subjected to the flow cytometry analysis of apoptosis. **(G)** The final results were shown by the column charts, which were representative of at least three independent experiments being each performed in triplicate. Of note, *$$, p <0*.*01*, and *$, p < 0*.*05* indicate significant increases in each basal value of [[*Nrf1α*^*–/–*^]_*T0*_, [*Nrf2*^*–/–△TA*^]_*T0*_, and [*caNrf2*^*ΔN*^]_*T0*_ with that of [*WT*]_*T0*_, while both *\* p < 0*.*05* and *\*\* p < 0*.*01* denote significant decreases of those values from each of cell lines treated by DTT for 4 h (i.e., [X]_*T4*_) or 24 h (i.e., [X]_*T24*_) versus its untreated [X]_*T0*_ value in the same groups.

Further experiments revealed that an intrinsic significant augmentation in the proportion of GSSG to GSH was *de facto* in *Nrf1α*^*–/–*^ cells, whereas *Nrf2*^*–/–ΔTA*^ cells only had a modestly enhanced ratio of GSSG to GSH, when compared with the *WT*_*t0*_ control (Fig. 4E). Upon stimulation of DTT for 4 h, an inducible elevated rate of GSSG to GSH was examined only in *WT* and *Nrf2*^*–/–ΔTA*^, but not *Nrf1α*^*–/–*^ *or caNrf2*^*ΔN*^ cells, which seemed closely to be the highly endogenous level obtained from *Nrf1α*^*–/–*^ cells. When stimulation of DTT extended to 24 h, such DTT-stimulated elevation descended closely to their basal levels (Fig. 4E). Notably, DTT intervention of *Nrf1α*^*–/–*^ cells for 4 h to 24 h resulted in substantial decreased rates of GSSG to GSH, but no changes in the GSSG to GSH ratio was observed in *caNrf2*^*ΔN*^ cells. From these, it is inferable to be attributable to aberrant accumulation of hyperactive Nrf2 in *Nrf1α*^*–/–*^ cells insomuch as to reinforce its antioxidant and detoxifying cytoprotection against DTT, whereas *caNrf2*^*ΔN*^ cells with a genetic deletion of the N-terminal Keap1-binding domain has lost its powerful response to the redox-sensing by Keap1.

Further experimental evidence was also obtained from flow-cytometry analysis of cell apoptosis, revealing a lowest number of apoptosis of untreated *WT* cells, while a relative higher number of *Nrf1α*^*–/–*^ cells underwent apoptosis (Fig. 4, F & G). Almost similar apoptosis of *Nrf2*^*–/–ΔTA*^ and *caNrf2*^*ΔN*^ cells was closely to that of *WT* cells, but much lower than that of *Nrf1α*^*–/–*^ cells (Fig. 4F). However, DTT stimulation caused significant decreases in apoptosis of *Nrf1α*^*–/–*^ cells, but slightly increased apoptosis of *Nrf2*^*–/–ΔTA*^ or *caNrf2*^*ΔN*^ cells at 24 h after this chemical treatment, along with no changes in *caNrf2*^*ΔN*^ cells (Fig. 4G). This demonstrate that accumulated Nrf2 in *Nrf1α*^*–/–*^ cells has still exerted its intrinsic cytoprotective effect against DTT-induced apoptosis, but this effect is lost in both *Nrf2*^*–/–ΔTA*^ and *caNrf2*^*ΔN*^ cell lines (the former *Nrf2*^*–/–ΔTA*^ lacks its transactivation domain to regulate its target genes, while the latter *caNrf2*^*ΔN*^ lacks its responsive interaction with the redox-sensing Keap1).

### 3.5 The redox status of Cys342 and Cys640 in Nrf1 is required for its protein stability and *trans*-activity

Our previous studies showed that Nrf1 undergoes a variety of post-translational modifications, such as glycosylation, deglycosylation, ubiquitination and phosphorylation, as well selectively proteolytic processing (12). Here, we investigate which cysteine (Cys) residues within Nrf1 can directly sense the reductive stressor DTT (that can reduce protein disulfide bond (-*S*-*S*-) to sulfhydryl (-*SH*) and also oxidize itself into six-membered ring, to assist in maintaining protein function (33)). A schematic shows mutagenesis mapping of four Cys residues of Nrf1 into serines (i.e., C342S, C521S, C533S and C640S) within its NST, Neh6L and bZIP domains, respectively (Fig. 5A). Next, the pulse-chase experiments revealed that a considerable portion of the full-length glycoproteins of two mutants Nrf1^C342S^ and Nrf1^C640S^ were rapidly converted into their deglycoproteins and then proteolytically processed to disappear in a faster manner than those equivalents arising from wild-type Nrf1 (Fig. 5B). These protein stability was evaluated by western blotting to measure their half-lives of Nrf1^C342S^, Nrf1^C640S^ and its wild-type Nrf1, which were calculated for their glycoprotein turnover to be 0.36 h (21.6 min), 0.35 h (21 min) and 1.27 h (76.2 min) (Fig. 5C), and for their deglycoprotein turnover to be 0.45 h (27 min), 0.66 h (39.6 min) and 1.58 h (94.8 min) (Fig. 5D), respectively, after treatment with CHX (to inhibit the nascent polypeptide synthesis). By contrast, the stability of another two mutants Nrf1^C521S^ and Nrf1^C533S^ were only marginally affected, when compared with that of wild-type Nrf1 (Fig. 5B to D).

**Figure 5.**
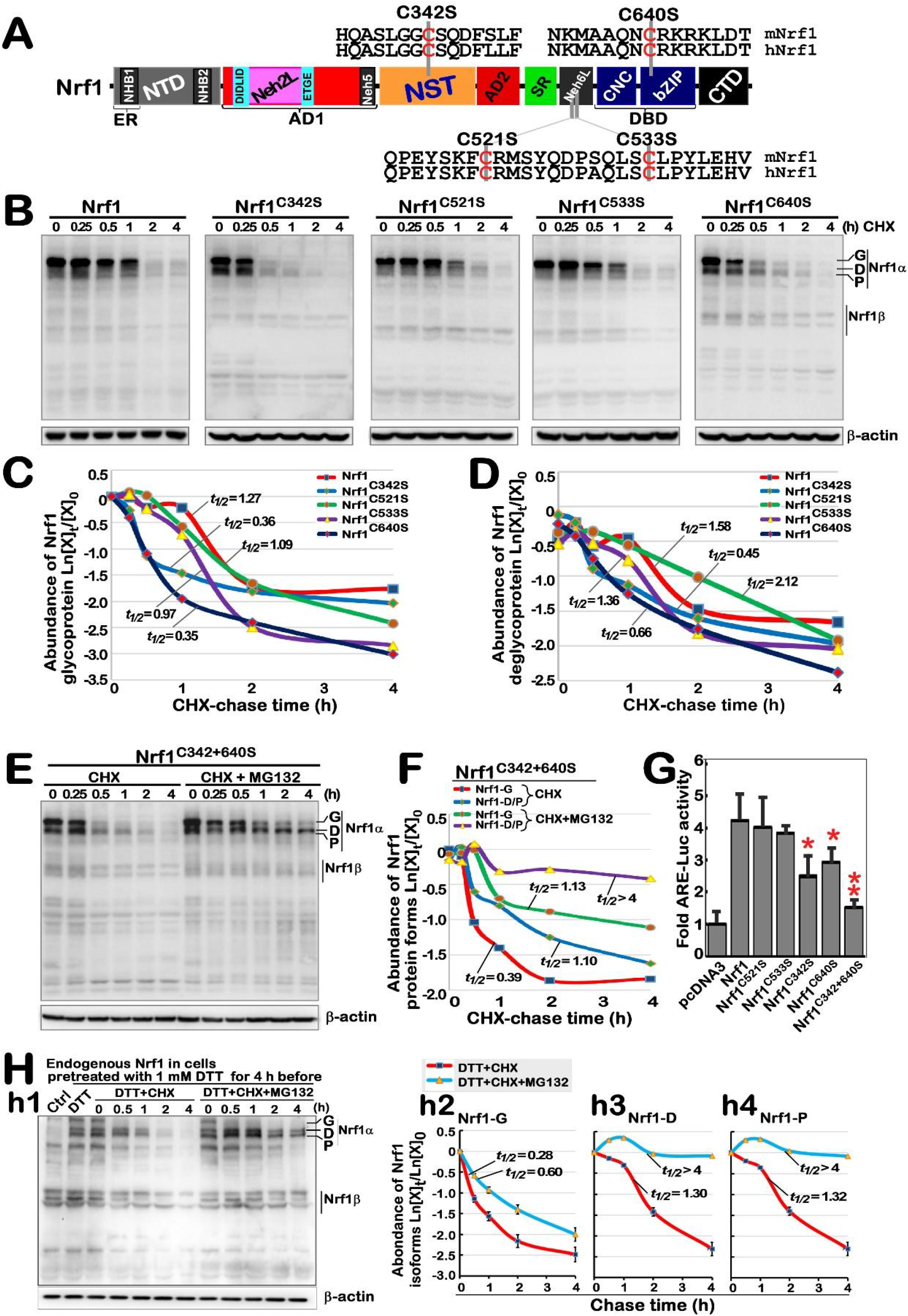
Nrf1 stability and its *trans*-activity were determined by redox status of its Cys342 and Cys640. **(A)** Schematic diagram of four cysteine mutants within Nrf1, which are distributed in its NST, Neh6L and bZIP domains. **(B)** Stability of Nrf1 and its mutants was determined by the pulse-chase experiments for 0 to 4 h intervention by CHX (5 μg/mL). **(C**,**D)** The results was evaluated for the turnover trend of glycoproteins (**C**) and deglycoproteins (including processed proteins) (**D**) of Nrf1α and mutants, with distinct half-lives estimated. **(E**,**F)** After treatment of cells with CHX alone or plus10 μM MG132, stability of the Nrf1^C342/640S^ mutant was further examined by pulse-chase experiments **(E)** and distinct half-lives of its derivative proteins were calculated **(F). (G)** *WT* cells were co-transfected with an *ARE-Luc* reporter, together with each of indicated expression constructs for Nrf1, its mutants or empty pcDNA3.1 vector. The luciferase activity was normalized to their internal controls and corresponding backgrounds obtained from the co-transfection of cells with non-*ARE* reporter and each of the expression constructs. The results were calculated as *mean ± SD* (*n= 6×3*) relative to the basal activity (at a given value of 1.0) obtained from the transfection of cells with empty pcDNA3.1 vector and ARE-driven reporter. **(H)** Stability of endogenous Nrf1α proteins was examined by pulse-chase experiments after stimulation of cells by 1 mM DTT (*h1*), and changes in its derived glycoprotein, deglycoprotein, processed protein abundances were represented by their respective turnover time-course curves (*h2, h3, h4*). In addition, similar examinations of ectopically-expressing Nrf1α and its Nrf1^C342/640S^ mutant after 1 mM DTT stimulation were also shown in **Figure S4**. The intensity of relevant immunoblots representing different protein expression levels was also quantified by the Quan-tity One 4.5.2 software. The resulting data were shown graphically, after being calculated by a formula of Ln ([X]_t_/[X]_0_), in which [X]_t_ indicated a fold change (*mean ± SD*) in each of those examined protein expression levels at different times relative to corresponding controls measured at 0 h (*i*.*e*., [A]_0_), which were representative of at least three independent experiments.

Further examination of a bi-Cys mutant Nrf1^C342/640S^ unraveled that half-lives of its glycoprotein and deglycoprotein turnover were calculated to be 0.39 h (23.4 min) and 1.10 h (66.0 min), respectively, after CHX treatment (Fig. 5E, 5F). Upon addition of proteasomal inhibitor MG132 plus CHX, the half-life of this mutant glycoprotein was only modestly extended to 1.13 h (67.8 min), just because it has to be deglycosylated during the pulse-chase experiments. Rather, the half-life of this mutant deglycoprotein was significantly extended to over 4 h before stopping this experiments (Fig. 5E, 5F). This implies that this deglycoprotein turnover is quality-controlled by proteasome-mediated degradation pathway. Interestingly, transactivation activity of *ARE*-driven luciferase reporter gene regulated by Nrf1 was significantly inhibited by its mutants Nrf1^C342S^, Nrf1^C640S^ or Nrf1^C342/640S^ (Fig. 5G). Thereby, it is inferable that the redox state of both Cys342 and Cys640 residues in Nrf1 could be required for its protein stability and transcriptional activity.

Next, the effect of DTT-induced redox stress on Nrf1 stability was further determined. The results showed that DTT enhanced accumulation of all the endogenous Nrf1-drived proteins, and they were further accumulated by MG132 plus DTT (Fig. 5H, *h1*). However, the half-lives of Nrf1 glycoprotein (i.e., Nrf1-G), deglycoprotein (i.e., Nrf1-D) and its processed protein (i.e., Nrf1-P) were measured to be 0.28 h (16.8 min), 1.30 h (78 min), and 1.32 h (79.2 min), respectively, after DTT co-treatment of cells with CHX (Fig. 5H, *h2* to *h4*). Upon addition of MG132 to DTT/CHX-treated cells, the half-life of Nrf1-G was slightly extended to 0.60 h (36 min), whereas the half-lives of Nrf1-D and Nrf1-P were strikingly prolonged to over than 4 h after stopping experiments. Further determination of Nrf1 protein turnover was carried out by Western blotting of ectopically-expressed isoforms of this CNC-bZIP factor and its bi-Cys mutant Nrf1^C342/640S^, which were resolved by gradient LDS-NuPAGE gels containing 4%-12% polyacrylamides (Fig. S4). The results revealed that abundance of Nrf1 was enhanced by DTT (Fig. S4A). In the presence of DTT, the half-lives of Nrf1-G, -D and -P were calculated to be 0.18 h (10.8 min), 1.06 h (63.6 min) and 0.74 h (44.4 min), respectively, after CHX treatment (Fig. S4A, *a2* to *a4*). By contrast, Nrf1^C342/640S^ appeared to be unaffected by this chemical (Fig. S4B, *b1*), with distinct half-lives of its isoforms that were slightly changed to be 0.24 h (14.4 min), 0.64 h (38.4 min) and 0.57 h (34.2 min), respectively after co-treatment of DTT with CHX (*b2 to b4*). Notably, the glycoprotein half-life of Nrf1 or Nrf1^C342/640S^ was only modestly extended by addition of MG132 to be 0.30 h (18 min) or 0.50 h (30 min), respectively, but their deglycoproteins and proteolytic proteins were all significantly prolonged to over than 4 h. Taken together, these data indicate that the redox state of Cys342 and Cys640 residues in Nrf1 is not only required for its protein stability and both may also serve as a redox sensor for the DTT stressor.

### 3.6 Biphasic effects of DTT on transcriptional expression of Nrf1-target proteasomal genes

It was previously reported that Nrf1, rather than Nrf2, plays an essential role in controlling transcriptional expression of all proteasomal subunit genes (34). Such expression of the proteasomal genes regulated by Nrf1 is required for the ER-associated degradation (ERAD), as accompanied by induction of three classical response pathways driven by the ER stress-sensing genes (i.e., *PERK, IRE1* and *ATF6*). Thereby, we here explored whether DTT-stimulated Nrf1 and Nrf2 is also required for the expression of key proteasomal (*PSM*) genes along with ER signaling networks. The RT-qPCR results showed significant decreases in basal mRNA expression levels of all three examined genes *PSMB5, PSMB6* and *PSMB7* in *Nrf1α*^*–/–*^ cells (Fig. 6A), while basal *PSMB5* and *PSMB7* expression levels were also partially decreased in *Nrf2*^*–/–ΔTA*^ or *caNrf2*^*ΔN*^ cells, when compared with *WT*_*t0*_ controls. This implies that except for Nrf1, Nrf2 is also partially involved in regulating transcription of some PSM genes (e.g., *PSMB5* and *PSMB7*) through its N-terminal Keap1-binding domain, in addition to its transactivation domain.

**Figure 6.**
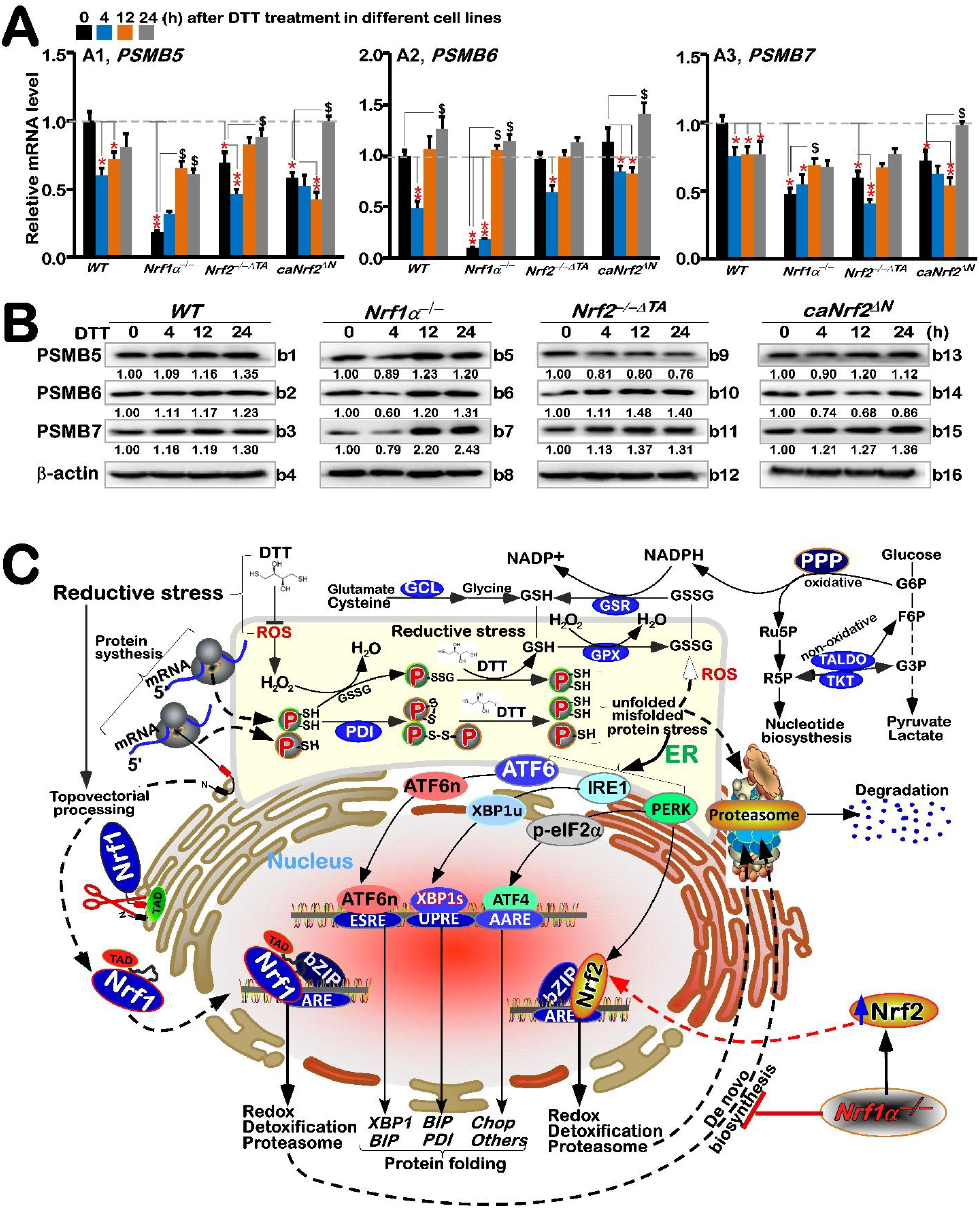
Differential expression of proteasomal genes regulated by Nrf1-independent pathway under reductive stress induced by DTT. **(A)** Four genotypic cell lines of *WT, Nrf1α*^*–/–*^, *Nrf2*^*–/–△TA*^ and *caNrf2*^*△N*^ were (or were not) treated with 1 mM DTT for 0 to 24 h, before both basal and DTT-inducible mRNA expression levels of core proteasomal subunit genes (i.e., *PSMB5* (*A1*), *PSMB6* (*A2*) and *PSMB7* (*A3*) were determined by RT-qPCR. The resulting data were shown as fold changes (*mean ± SD, n = 3 × 3*), which are representative of at least three independent experiments being each performed in triplicates. Significant increases (*$, p < 0*.*05; $$, p < 0*.*01*) and significant decreases (*\* p < 0*.*05; ** p < 0*.*01*) were statistically calculated by comparison with their corresponding controls (measured at 0 h). **(B)** These proteasomal protein expression levels in four distinct cell lines were determined by western blotting with their indicated antibodies, whilst *β*-actin served as a loading control. The intensity of those immunoblots, representing different protein expression levels, was also quantified by the Quantity One 4.5.2 software as shown on *bottom* **(C)** A proposed model is to provide a better explanation of distinct roles for Nrf1 and Nrf2 in DTT-stimulated reductive stress response, along with ER stress signaling and relevant redox metabolism.

Interestingly, DTT stimulation of *Nrf1α*^*–/–*^ cells (with accumulation of hyperactive Nrf2) caused evident increases in *PSMB5, PSMB6* and *PSMB7* at their mRNA and protein expression levels closely to basal *WT*_*t0*_ controls (Fig. 6A and 6B, *b5* to *b7*). This indicates such up-regulation of proteasomal genes by DTT may occur through an Nrf1-independent and/or Nrf2-dependent pathways. In *WT* cells, DTT caused partial decreases in *PSMB5* and *PSMB7*, and also biphasic changes (i.e., early decreased, then recovered and even elevated) in *PSMB6* (Fig. 6A), but their protein abundances were almost unaltered (Fig. 6B, *b1* to *b3*). Similarly, DTT-inducible biphasic expression of all three examined genes were also observed in *Nrf2*^*–/–ΔTA*^ or *caNrf2*^*ΔN*^ cell lines, with no obvious changes in their protein levels. Together, these suggest transcriptional expression of proteasomal genes is also tightly regulated by other not-yet-identified factors (e.g. Bach1), along with Nrf1 and Nrf2, particularly under DTT-stimulated stress conditions, albeit the basal proteasomal expression is predominantly governed by Nrf1, as well by partially Nrf2.

## 4. Discussion

The concept of ‘Stress’ was first put forward by Selye H, a Canadian physiologist, which represents a state of tension when an individual feels threatened physically and mentally (35). Similar concept of stress in cell biology is also employed, and the relevant topic has become an attractive direction to be followed by a large number of researchers from different fields. Most studies on animal stress are primarily focused on cell physiological basis, so mechanistic studies on cell stress have provided part of theoretical basis for animal stress. The occurrence of cell stress is to enhance the ability of cells to resist damage and thus survive under adverse conditions. Such results are caused by distinct stressors, including external chemical or physical stimuli (such as drugs, pathogens, or radiation) and internal factors (such as nutritional deficiencies). According to the classification of stress-generating conditions, different types of cell stresses are divided into redox (i.e., oxidative and reductive) stress, heat stress, hypoxia stress, genotoxic stress and nutritional stress. By search of oxidative stress from the PubMed database, there are 280,792 entities reported until May 24, 2022; and this number is also rapidly increasing yearly. This indicates the importance of oxidative stress attracting a great deal of attentions from life sciences, medicines and other relevant areas. Of note, ROS were mainly derived from the mitochondria, macrophages and non-macrophage’s nicotinamide adenine dinucleotide phosphate (NADPH) oxidase system, neutrophils, xanthine oxidase system and hypoxanthine oxidase system. One of the main mechanisms of tissue injury is predominantly triggered by distinct pathological processes, such as tumor, diabetes, atherosclerosis, fatty liver and organ ischemia-reperfusion (I/R) injury. Interestingly, there is a close relationship between oxidative stress and ER stress, because oxidative stress induced by pathological stimuli can also provoke the occurrence of ER stress, and vice versa. This implies there exists a mutually promoting positive feedback circuit between both oxidative and ER stresses (36).

As a matter of fact, the ER lumen is known as the most oxidizing subcellular compartment in cells, which facilitates formation of disulfide bonds in relevant protein structures and their proper folding. This process is also accompanied by ROS to be released under ER stress, albeit the mitochondria are viewed as a key site for ROS byproducts. Such oxidative circumstances are provided for synthesis of biological macromolecules in the ER. In the meantime, appropriate oxidative stress triggers a certain hormestic effect to assist in the reprocessing of unfolded and misfolded proteins in the ER. Rather, the excessive ROS cause ER stress and result in accumulation of unfolded and misfolded proteins. Such UPR effect can also be triggered by the functional load of ER. Under normal conditions, the ER stress sensors IRE1, PERK, ATF6 bind to GRP78 in the ER and also remain inactive. Upon dissociation of GRP78 from the ER lumen, it leads to the oligomerization of transmembrane protein IRE1, transfer of ATF6 to the Golgi apparatus, and activation of PERK, respectively (Fig. 6C). The latter PERK phosphorylates and activates eIF2α so as to terminate to the general translation process, thus reducing the internal load of ER. The active phosphorylated IRE1 splices downstream *XBP1* mRNA to give rise to the transcriptional activity of XBP1s (i.e., spliced *XBP1* encodes a longer polypeptide than its unspliced prototype *XBP1u* (with resistance to *XBP1s’* function), which enters the nucleus and continuously activates UPR genes by binding the ER stress elements (ERSEs) in their promotor regions (37) (38). These indicate a complex and large regulatory relationship network between the UPR to ER stress and the occurrence of oxidative stress. In UPR response, CHOP acts as a pro-apoptotic factor to mediate the activation of ER stress-related apoptotic pathways. To combat overproduction of ROS, almost all cellular life forms have evolutionarily been equipped with an array of naturally powerful antioxidant defenses. These include a series of essential antioxidant, detoxification and cytoprotective mechanisms controlled by the CNC-bZIP transcription factors (39). In this family, Nrf1 can activate transcriptional expression of Herpud1 (homocysteine-inducible ER-resident protein with ubiquitin-like domain 1, also called Herp1, that is involved in both UPR and ERAD) driven by its ARE battery, and the resulting ubiquitin complex is transferred to the proteasome in order to promote the unfolded and misfolded protein degradation, thereby preventing or alleviating the ER-overloaded stress (40). This notion is supported by experimental evidence that endogenous ER stress signaling to activate UPR occur in the steatotic hepatocytes cells with homozygous knockout of *Nrf1*^*–/–*^, but not of *Nrf2*^*–/–*^(21). Therefore, it is inferable that Nrf1 plays an important role in maintaining the ER redox homeostasis, particularly upon sensing intracellular protein, lipid, glucose and redox challenges. The loss of this function in mice results in overt severe pathological consequence, that is caused through *Nrf1α*^*–/–*^ cell proliferation and malignant transformation, leading to spontaneous development of non-alcoholic steatohepatitis and hepatocellular carcinoma (41).

In stinking contrast with oxidative stress, only a small number of 7,864 entities about reductive stress are searched from the PubMed database until May 24, 2022. In this study, we examine distinct responsive effects of Nrf1 and Nrf2 to DTT-stimulated reductive stress. This is owing to the inherent reducibility of DTT to maintain the reduced state of protein sulfhydryl groups, which is crucial important for the stability of many protein functions. However, protein disulfide bond (-*S*-*S*-) was reduced by DTT into sulfhydryl (-*SH*), which affects proper folding of many proteins and even triggers ER stress induced by misfolded and unfolded proteins (42). This is because the unstable cysteine (Cys) is the only amino acid with reducing sulfhydryl (-*SH*) in more than 20 amino acids that make up the proteins, and thus is prone to oxidation-reduction to be converted with the -S-S-bond of cystine. Herein, we found that the redox status of Cys342 (in Nrf1’s glycodomain) and Cys640 (in Nrf1’s DNA-binding domain) residues is required for this CNC-bZIP protein stability and trans-activity.

Nrf1 is a *de facto* mobile transmembrane protein with a dynamic membrane topology, which is somewhat similar to, but different from, that of PERK, IRE1 and ATF6 within and around the ER (19). Here, our evidence suggests that PERK, IRE1, and ATF6 are differentially activated along with differential expression of their cognate responsive genes by DTT treatment of different genotypic cell lines with the presence or absence of Nrf1 and Nrf2 (for different time-dependent effects). Specifically, DTT-induced ERS response signaling pathway is accompanied by the transcriptional expression of Nrf1 and Nrf2. The bidirectional expression of PERK and eIF2α, that are positively and negatively regulated by Nrf1 and Nrf2, respectively, is accompanied by down-regulation of ATF4 by DTT, albeit the latter basal expression was upregulated by *Nrf1α*^*–/–*^ and *caNrf2*^*ΔN*^. The IRE1-XBP1 and ATF6 were differentially up-regulated by DTT in *Nrf1α*^*–/–*^ and *Nrf2*^*–/–ΔTA*^, but XBP1 appeared to be unaffected by *caNrf2*^*ΔN*^. These indicate that DTT-caused reductive stress leads to trans-repression of ATF4, but transactivation of XBP1 and ATF6, possibly dependent on Nrf1 and Nrf2; the latter N-terminal Keap1-binding domain is also required for this responsiveness to DTT. The UPR signaling-provoked chaperone GRP78 (also called BIP) was rapidly induced by DTT, and then gradually decreased and even suppressed by this reductive stressor, whereas PDI (catalyzing disulfide bonds to be formed in proper folding proteins) was gradually upregulated by DTT as this treatment time was extended, in all those examined cell lines, regardless of whether Nrf1 is present or Nrf2 lost either its N-terminal Keap1-binding domain or transactivation domain. This demonstrates that DTT cannot only rapidly stimulate ER stress signaling and also provoke a rebound effect on PDI to increment its expression levels, but this process has not to rely on Nrf1 or Nrf2, albeit both factors may be involved in.

Upon stimulation of wild-type cells by DTT, those ARE-driven genes encoding GCLM, GCLC, HO-1, NQO1, GSR, TKT, and TALDO were up-regulated to different extents. Among them, basal and DTT-stimulated expression levels of GCLM, GCLC, HO-1 and TKT were further augmented in *Nrf1α*^*–/–*^ cells, but obviously suppressed in *Nrf2*^*–/–ΔTA*^ cells, and almost unaffected in *caNrf2*^*ΔN*^ cells except from only lagged induction of GCLC. This implies they are Nrf2-dependent responsive genes to DTT, and are also regulated by its N-terminal Keap1-binding domain. By contrast, DTT-inducible expression of GSR, GPX1 and NQO1 were roughly unaffected by *Nrf1α*^*–/–*^ and *Nrf2*^*–/–ΔTA*^, suggesting they are co-regulated by Nrf1 and Nrf2. Only marginally lagged stimulation of MT1E and MT2 by DTT were examined in wild-type cells, but totally abashed by knockout of *Nrf1α*^*–/–*^ or *Nrf2*^*–/–ΔTA*^, respectively. Conversely, the formal MT1E’s basal and DTT-stimulated expression levels were significantly increased in *Nrf2*^*–/–ΔTA*^ cells, while the latter MT2 expression was modestly elevated in *Nrf1α*^*–/–*^ cells, which occurred after 16 h treatment of DTT, but its early treatment for 4 h to 12 h led to an overt inhibition. These indicate that MT1E is one of Nrf1-specific targets, whereas MT2 is another Nrf2-specific target, but both are not sensitive to DTT-leading reductive, particularly during early treatment, whereas the lagged induction may be triggered by other not-yet-unidentified mechanisms (in which an increase of ROS cannot be ruled out). Unexpectedly a marginal increase in ROS levels were determined only for 4 h intervention of *WT* or *Nrf1α*^*–/–*^ cell lines. The increased ratio of GSSG to GSH rate was increased by DTT in *WT* and *Nrf2*^*–/–ΔTA*^ cell lines, but repressed in *Nrf1α*^*–/–*^ (with hyperactive Nrf2 accumulated) or almost unaffected by *caNrf2*^*ΔN*^. This suggests DTT may induce reductive stress through Nrf2 to yield a certain amount of GSH in *Nrf1α*^*–/–*^ cells, which enables cytoprotection against this chemical cytotoxicity, whereas loss of *Nrf2*^*–/–ΔTA*^ also leads to mild oxidative stress triggered by DTT, but detailed mechanisms remain to be explored in the future work.

## 5. Concluding remarks

Hitherto, the overwhelming majority of redox researches are focused principally on oxidative stress and antioxidant responses, owing to the relative rarity of workers on reductive stress and anti-reductant response. Herein, we found that DTT, as reductive stressor, stimulates distinct expression profiles of Nrf1 and Nrf2, along with their cognate target genes driven by antioxidant and/or electrophile response elements (AREs/EpREs). Their differential expression levels are also affected by the self-recovery (protective) ability of stimulated cells against such reductive drug cytotoxicity. By analyzing distinct expression levels of each gene in different genetic backgrounds of *WT, Nrf1α*^*–/–*^, *Nrf2*^*–/–ΔTA*^ and *caNrf2*^*ΔN*^ cell lines, it was confirmed distinct roles of Nrf1 and Nrf2 in mediating differential expression levels of both ARE-driven and UPR-target genes (Figure 6C). For examples, XBP1 and Nrf1 are closely related, while HO-1, IRE1, CHOP and ATF4 genes were closely related to Nrf2. Albeit Nrf2 can serve as an upstream transcriptional regulator of Nrf1, it is inferred, according to the expression situation under this drug induction, that Nrf1 mainly regulates the expression of antioxidant and ER stress genes under normal state of cells, while Nrf2 mainly regulates the expression of intracellular genes under induced stress state. Overall, Nrf1 and Nrf2 are required for coordinated regulation of DTT-leading reductive stress response. However, DTT-stimulated expression of Nrf1-target proteasomal genes was still detected in *Nrf1α*^*–/–*^ cells, but roughly unaffected by constitutive activation of Nrf2 (i.e., *caNrf2*^*ΔN*^) or its inactivation (i.e., *Nrf2*^*–/–ΔTA*^), demonstrating a requirement for such DTT-induced proteasomal genes regulated by an Nrf1/2-indepdent pathway.

## Supplementary Materials

The following are available materials, **Figure S1**: The protein expression of Nrf1 and Nrf2 under the reductive stress induced by DTT; **Figure S2**: The expression level of redox response proteins induced by DTT in different genotypes; **Figure S3**: The expression level of ER stress related proteins induced by DTT in distinct cells; **Figure S4**: The half-lives changing of Nrf1-G, -D and -P (endogenous and exogenous) after treated by DTT.

## Author Contributions

Both R.W., Z.F. and J.Y. performed the experiments with help of Z.Z. and S.H., collected all the relevant data, and wrote a draft of this manuscript with most figures and supplemental information. Y.Z. designed and supervised this study, analyzed all the data, helped to prepare all figures with cartoons, and wrote and revised the paper with help of G.S. All authors have read and agreed to the published version of the manuscript.

## Institutional Review Board Statement

Not applicable in this study without human or animal.

## Informed Consent Statement

Not applicable in this study without human experiments.

## Data Availability Statement

All data needed to evaluate the conclusions in the paper are present in this publication along with the supplementary information documents that can be found online. The additional data related to this paper may also be requested from the corresponding author on reasonable request.

## Acknowledgments

We are greatly thankful to Lu Qiu (Zhengzhou University, Henan, China) and Yonggang Ren (North Sichuan Medical College, Sichuan, China) for their involvement in establishing the indicated cell lines used in this study. We also thank all other present and past members of Zhang’s laboratory (at Chongqing University, China) for giving critical discussion and invaluable help with this work.

