## Supplement figure S1-S4 for "Distinct roles of Nrf1 and Nrf2 in monitoring the reductive stress response to dithiothreitol (DTT)"

#### Supplemental materials:

### Figure S1

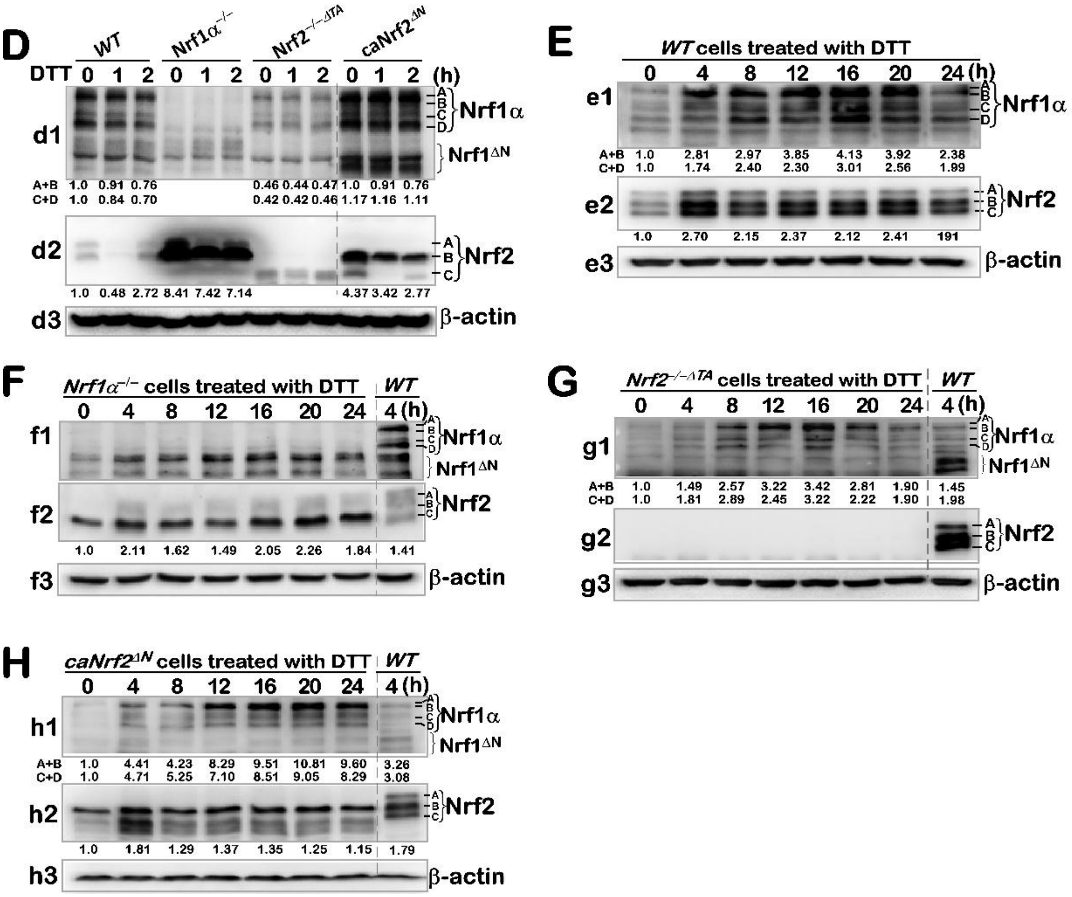


**Figure S1**. The protein expression of Nrf1 and Nrf2 under the reductive stress induced by DTT.

Figure. S2

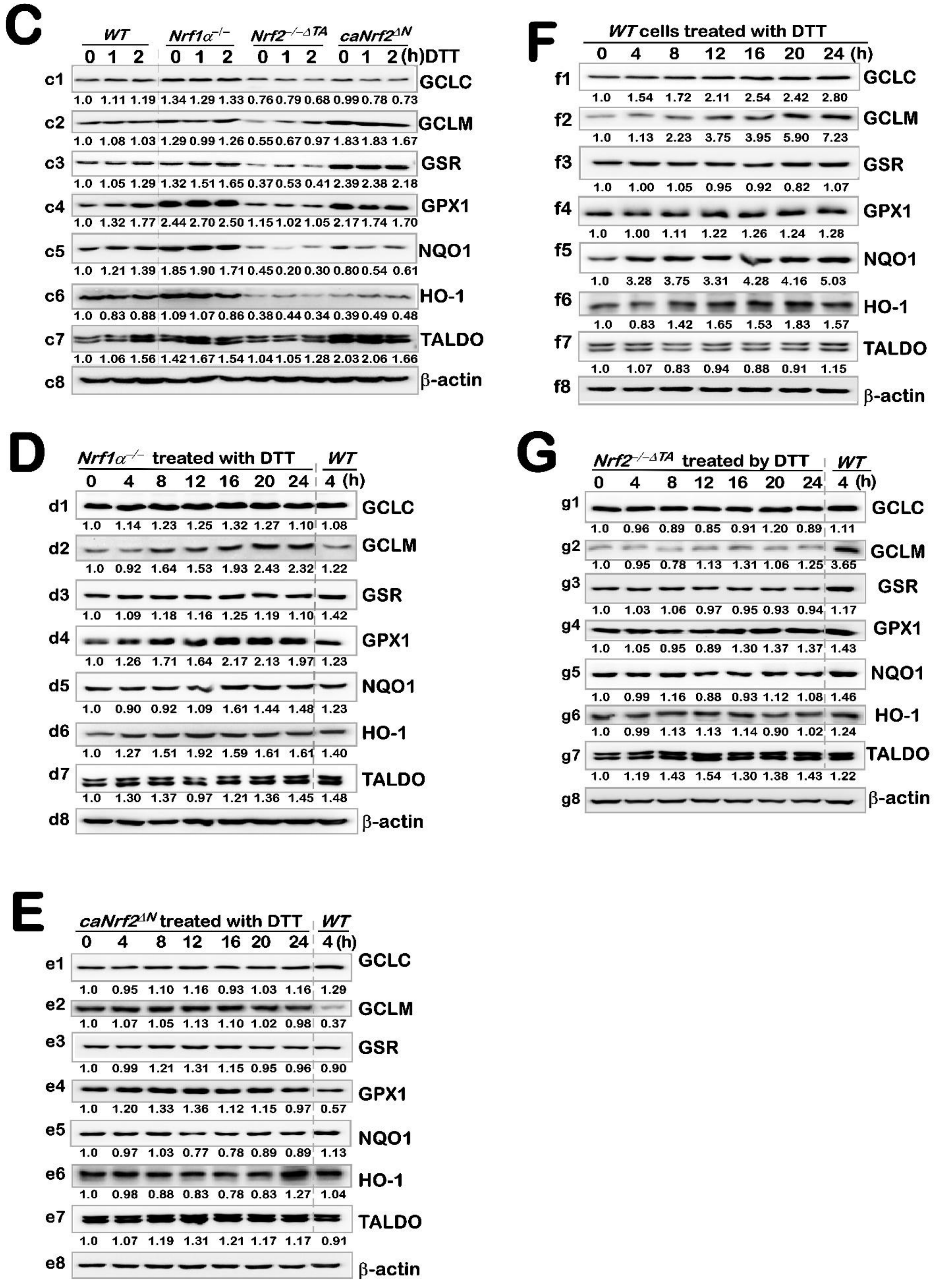


**Figure S2**. The expression level of redox response proteins induced by DTT in different genotypes.

### Figure S3

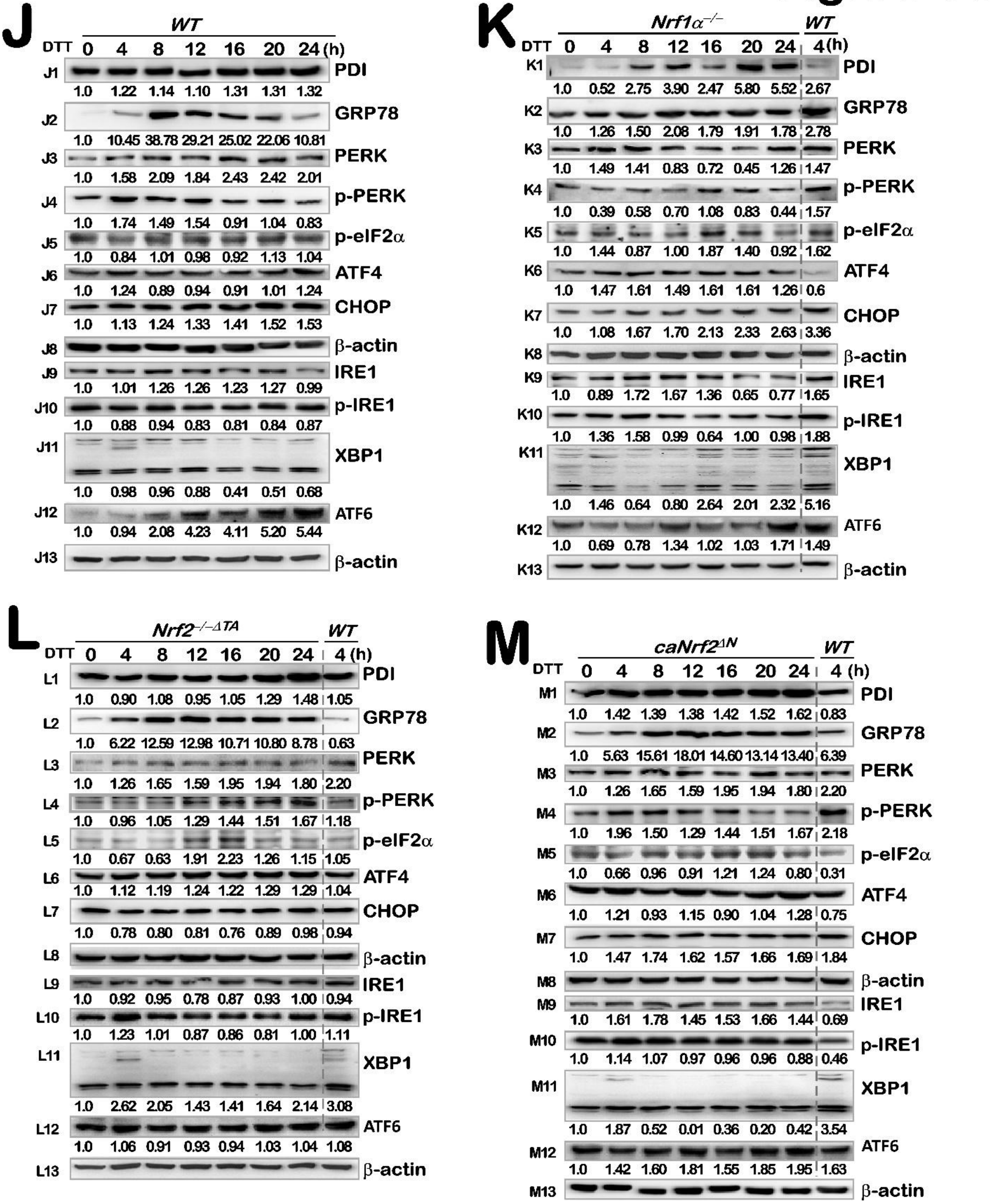


**Figure S3**. The expression level of ER stress related proteins induced by DTT in distinct cells.

### Figure S4

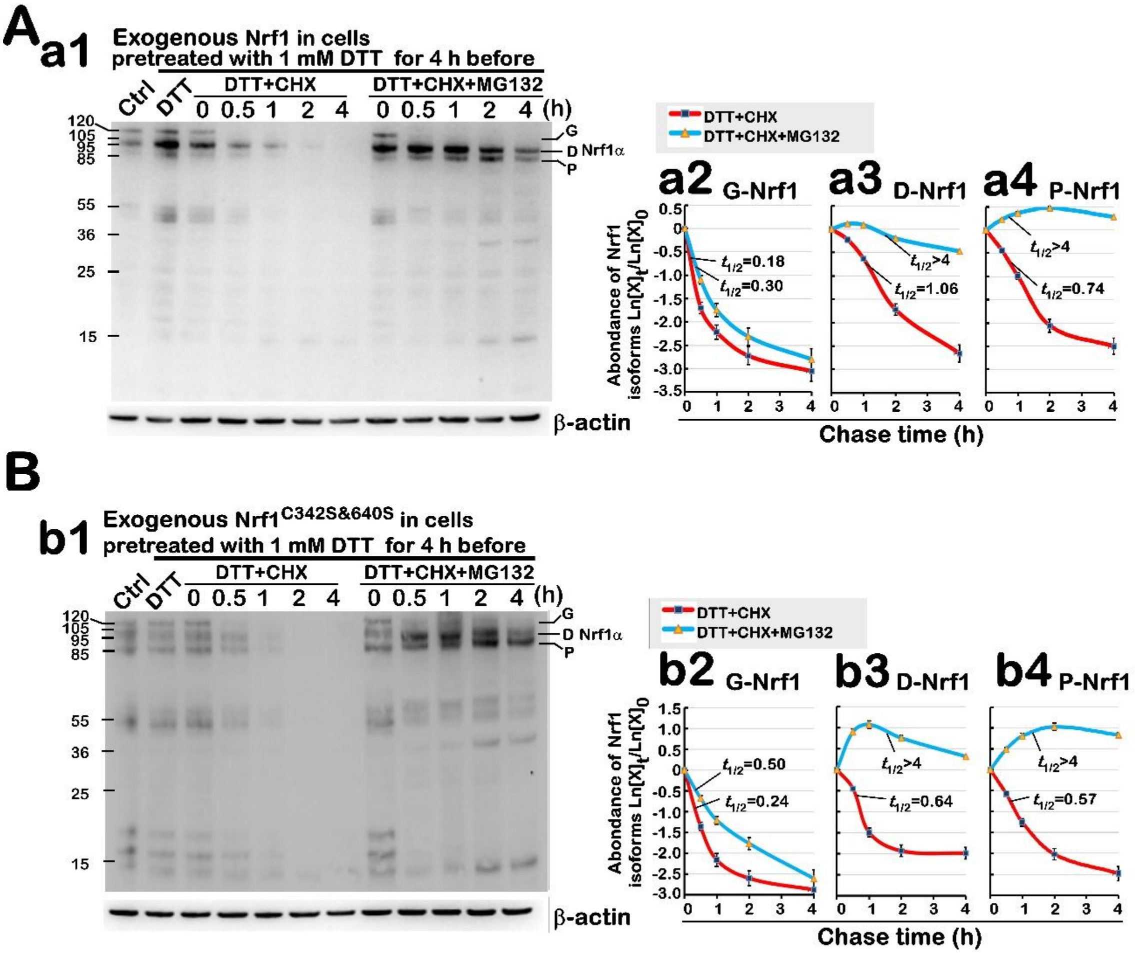


**Figure S4**. The half-lives changing of Nrf1-G, -D and -P (endogenous and exogenous) after treated by DTT.
